# The phylogeny of Triticeae Dumort. (Poaceae): resolution and phylogenetic conflict based on a genome-wide selection of nuclear loci

**DOI:** 10.1101/2024.05.22.595384

**Authors:** Roberta J. Mason-Gamer, Dawson M. White

## Abstract

**Premise:** The wheat tribe, Triticeae, has been the subject of molecular phylogenetic analyses for nearly three decades, and extensive phylogenetic conflict has been apparent from the earliest comparisons among DNA-based data sets. While most previous analyses focused primarily on nuclear vs. chloroplast DNA conflict, the present analysis provides a broader picture of conflict among nuclear loci throughout the tribe.

**Methods:** Exon data were generated from over 1000 nuclear loci using targeted sequence capture with custom baits, and nearly-complete chloroplast genome sequences were recovered. Phylogenetic conflict was assessed among the trees from the chloroplast genomes, the concatenated nuclear loci, and a series of nuclear-locus subsets guided by *Hordeum* chromosome gene maps.

**Key results:** At the intergeneric level, the analyses collectively reveal a few broadly consistent relationships. However, the prevailing pattern is one of extensive phylogenetic conflict throughout the tribe, among both deep and shallow branches, and with the extent of the conflict varying among data subsets.

**Conclusions:** The results suggest continual introgression or lineage sorting within and among the named lineages of the Triticeae, shaping both deep and shallow relationships in the tribe.

## INTRODUCTION

The wheat tribe, Triticeae, is a small group—its ∼500 species comprise only about 4% of the grass family (Soreng et al., 2015)—but the attention paid to its evolutionary history has been disproportionate to its size. This is primarily due to wide interest in its economically important species, which include wheat, barley, and rye, along with a number of agriculturally significant weeds (Swearington and Bargeron, 2016), and useful forage and range species (Asay et al., 2001; Vogel and Jensen, 2001). However, seemingly straightforward questions about relationships among Triticeae species have remained stubbornly intractable through decades of studies employing ever-advancing methods of data collection and analysis. As a result, generic classification within the tribe has changed dramatically through time (Barkworth, 2000; Bernhardt, 2015), and multiple conflicting taxonomic systems remain in simultaneous use.

Widespread allopolyploidy, ranging from tetraploid to at least dodecaploid genomes, complicates our estimations of Triticeae phylogeny. Polyploids unite closely- and/or distantly-related Triticeae genomes into numerous distinct combinations, some of which likely arose multiple times. Thus, in hindsight, it is unsurprising that some of the first cladistic analyses of morphological characters, which included both diploid and polyploid taxa (e.g., Baum, Estes, and Gupta, 1987), failed to resolve relationships. Later analyses that explicitly excluded polyploids (Kellogg, 1989; Seberg and Frederiksen, 2001) yielded fewer most-parsimonious trees and thus appeared to be more internally consistent, yet they were not in agreement with the few molecular phylogenies available at the time (Seberg and Frederiksen, 2001).

Cytogenetic data have been extensively applied to delimiting groups within the Triticeae, and these form the basis of what is referred to as the genomic classification of the tribe (Dewey, 1984; Löve, 1984). In the genomic system, delimitation of genera is based on the ability of chromosomes to pair at meiosis in interspecific hybrids, under the assumption that the degree of pairing reflects overall relatedness among genomes. Such analyses have also provided valuable information about the genomic complement of many polyploids, through assessment of meiotic chromosome pairing in hybrids between polyploids and potential diploid genome donors. However, genomic classification has been the subject of considerable skepticism (Baum, Estes, and Gupta, 1987; Kellogg, 1989; Seberg and Petersen, 1998). Furthermore, because pairing data are open to interpretation, there are differences among genome-based classification schemes even among workers who agree with the general approach. That said, morphological studies (e.g., Barkworth and Dewey, 1985; Barkworth et al., 2009) and molecular phylogenetic analyses (e.g., Mason-Gamer, 2013) at least partially support the genus-level groups that are defined by chromosome pairing data, and taxonomic classifications of the Triticeae based on cytogenetic data now prevail (Barkworth, 2005; Wang and Lu, 2014).

While chromosome-pairing analyses have contributed to the delimitation of genera, most of the recent studies of relationships *among* genera have applied molecular data. Chloroplast DNA (cpDNA) data sets have ranged from restriction sites (Mason-Gamer and Kellogg, 1996b), to sequenced loci (e.g., Petersen and Seberg, 1997; Mason-Gamer, Orme, and Anderson, 2002), to largely complete genomes (Bernhardt et al., 2017), and have converged on a generally well-supported phylogeny of the Triticeae chloroplast genomes. Other molecular analyses of the tribe primarily include DNA sequences from individual single- or low-copy genes (e.g., Mason-Gamer, 2001; Helfgott and Mason-Gamer, 2004; Mason-Gamer, 2005; Seberg and Petersen, 2007; Sun and Salomon, 2009; Sun and Zhang, 2011; Fan et al., 2013), highly-repetitive loci (Hsiao et al., 1995; Kellogg and Appels, 1995), and a multi-gene analysis of 1 cpDNA and 26 nuclear loci (Escobar et al., 2011). The results of these studies are difficult to generalize, due to a combination of lack of resolution in individual trees, and/or disagreement among them. None of the nuclear tree topologies to date closely resemble the cpDNA tree (Kellogg, Appels, and Mason-Gamer, 1996; Mason-Gamer and Kellogg, 1996a; Mason-Gamer, 2005; Seberg and Petersen, 2007).

The present analysis builds upon past molecular analyses of the wheat tribe through analyses of exon sequence data from over 1000 nuclear loci obtained from a broad sample of diploid representatives of the tribe using targeted sequence capture, along with a matched sample of nearly-complete chloroplast genomes. The results highlight (1) strongly-supported cpDNA-based relationships consistent with previous studies; (2) several well-supported “nuclear genome” relationships among major genera; (3) multiple lineages that remain difficult to place with confidence; (4) a few similarities and multiple broad differences between the chloroplast and combined-nuclear trees; and (5) variable relationships among nuclear data subsets, including conflicts with the combined-nuclear trees.

## MATERIAL AND METHODS

### Samples

The 76 DNA samples (Appendix S1; see Supplemental Data with this article) include 75 diploid accessions from the Triticeae (deposited at GH), most of which have been included in previous studies (e.g., Mason-Gamer and Kellogg, 1996a; Mason-Gamer, 2005, 2013). Of the multiple taxonomic systems in current use for the Triticeae, we generally follow Löve’s cytogenetic system of 37 genera, of which 13 comprise only allopolyploid species (not included here) and 24 comprise diploid/autopolyploid species. Some of Löve’s names have not entered common usage and are not used here, but our sample of 17 recognized genera includes at least one representative of 23 of his 24 diploid/autopolyploid cytogenetic genera— all except *Festucopsis* (Löve would classify our *Aegilops* sample into 6 genera; our *Thinopyrum elongatum* as *Lophopyrum elongatum*; and 9 of our 10 *Hordeum* species as *Critesion*). We include one outgroup, *Bromus tectorum*, representing the tribe Bromeae, which is sister to the Triticeae (e.g., Zhang et al., 2022). We initially included a representative of a second, more distant outgroup (*Avena fatua*, tribe Aveneae), but the overall sequence coverage was low, so this individual was excluded from the data analyses.

### Bait design, library preparation, target capture, and sequencing

Exon-specific baits were designed using the pipeline in Weitemier et al. (2014), which in brief: identifies coding sequences (exons) within the genome; eliminates loci with multiple, closely-related copies (paralogs); eliminates exons with high identities among different loci; filters loci according to desired minimum transcript length (960, in this case); and outputs a series of sequence “blocks” corresponding to exons, from which baits are designed. The first step of the pipeline employs BLAT (Kent, 2002), which uses transcriptome/cDNA sequences (without introns) to identify genomic coding sequences (including introns). This step was completed in iPlant (now Cyverse; http://www.cyverse.org/), and the remaining steps in the pipeline were carried out locally.

The pipeline used *Hordeum vulgare* (barley) cDNA sequences (Matsumoto et al., 2011) and a *Triticum urartu* (diploid wheat) transcriptome (Ling et al., 2013) to identify exons from the genomes of *H. vulgare* (International Barley Genome Sequencing Consortium, 2012) and *T. urartu* (Ling et al., 2013), respectively. Bait sequences were designed from the exon “blocks” from both *H. vulgare* and *T. urartu*. By most estimates, wheat and barley are distantly related within the tribe, so designing baits based on both was intended to facilitate sequence capture from across the tribe.

Loci with fewer than three exons (blocks) were deleted from the bait design set. The remaining loci were compared between wheat and barley using BLAST in Geneious v. 8.0 (Kearse et al., 2012). We retained loci with blocks of at least 80% similarity, to reduce the number of cases where the wheat and barley bait sets were targeting different genes. Finally, we used cd-hit-est-2D (Li and Godzik, 2006; Fu et al., 2012) to identify blocks with over 98% identity between wheat and barley, in which case the wheat copy was removed to prevent ordering redundant baits. Based on the final set of 18,048 blocks, 58,566 baits were designed and produced by MycroArray (now Daicel Arbor Biosciences, Ann Arbor, MI) using flexible spacing and 1X tiling.

The DNA samples were quantified with a Qubit fluorometer (Invitrogen), and libraries were prepared and sequenced in 3 sets at different times (“Library Set” column in Appendix S1) with 8, 32, and 40 individuals, respectively. For Set 1 only, each of the eight samples was used to prepare two duplicate libraries, so we could compare two sets of parameters in the later sequence capture protocols. For each DNA sample, 2.6 µg DNA was diluted in water to a volume of 130 µl, and sheared to ∼400 bp using a Covaris (Woburn, MA) M-220 focused-ultrasonicator at the Pritzker Lab, Field Museum of Natural History, following the parameters provided (50W peak incident power, 10% duty factor, 200 cycles per burst, 90 seconds, 20°C). Sheared samples were checked on 1% agarose gels. A volume of 50 µl (1 µg) of sheared DNA from each sample was used for library preparation using a KAPA Hyper Prep Illumina Kit (KAPA Biosystems, Wilmington, MA) and 48 barcoded adaptors (Bioo Scientific, Austin, TX) following the KAPA kit protocol. The ligation reactions were cleaned using Agencourt Ampure XP reagent (Beckman Coulter, Inc., Brea, CA) or KAPA Pure Beads (KAPA Biosystems) following the protocols supplied with each. Following cleaning, the prepared libraries were quantified with a Qubit fluorometer, and amplified 6 times (Sets 1 and 2) or 8 times (Set 3) with a KAPA Library Amplification Kit (KAPA Biosystems). The amplification products were cleaned using Agencourt Ampure XP reagent or KAPA Pure Beads, and quantified with a Qubit fluorometer. Prior to incubation with baits, the Set 1 samples from each of the 2 duplicate library preparations were combined to give 2 duplicate, 8-sample equimolar mixtures; the 32 samples from Set 2 were combined into 4 mixtures of eight samples each; and the 40 samples from Set 3 were combined into 8 mixtures of 5 samples. Each equimolar mixture had a total of 500 ng of DNA.

Targeted sequence capture and washing reactions for the Set 1 and 2 equimolar mixtures were carried out following the protocols in the MYbaits manual v. 2.3.1; for Set 3, we followed MYbaits manual v. 3.0.1. For Set 1, the two duplicate library mixtures were used to test two different sequence capture protocols: the first was incubated with the baits at 65°C for 36 hours and then washed at 65° (following the MyBaits v. 2.3.1 manual), and the other was incubated under less stringent conditions of 65°C for 12 hours, 62.5°C for 12 hours, and 60°C for 12 hours, and then washed at 60°C. We did not note any difference between the incubation protocols, so Sets 2 and 3 were incubated and washed at 65°C.

Post-capture libraries were amplified for 12 cycles following the MyBaits protocol. Amplified post-capture libraries were cleaned with Ampure XP bead-based reagent (Beckman Coulter, Inc.) and quantified on either an Agilent BioAnalyzer or Agilent TapeStation at the University of Illinois at Chicago DNA Services facility (UIC-DNAS). Post-capture library Set 1 was sequenced with a 160-basepair single-end run on one lane of an Illumina HiSeq 2500 at the Roy J. Carver Biotechnology Center at the University of Illinois at Urbana-Champaign. Sets 2 and 3 were sequenced at the UIC-DNAS on an Illumina NextSeq500 V2 with 150-bp paired-end reads: Set 2 was run on two mid-output lanes and Set 3 was run on a single high-output lane.

The post-hybridization washing protocol from the earlier MyBaits manual (v. 2.3.1) left enough off-target reads in Sets 1 and 2 to allow for cpDNA assembly from the post-hybridization reactions. The later MyBaits manual (v. 3.0.1, used for Set 3) was designed to be more efficient at removing off-target reads, and so it was; we did not obtain enough off-target chloroplast genome reads from Set 3 to assemble chloroplast sequences for these samples. Thus, 142 ng DNA (1µl) from each of the barcoded, pre-capture Set 3 library mixtures were pooled and sequenced at the UIC-DNAS in one mid-output lane on the Illumina NextSeq500 V2 with 150-bp paired-end reads. The chloroplast genome reads from both Set 3 runs were combined for cpDNA assembly.

### Data analysis

Raw reads were adaptor- and quality-trimmed using BBduk in Geneious 9.0 with the options kmer length=27, ktrim=right, minimum kmer=4, hamming distance=1, edit distance=0, quality trim=right/left, trim quality=6, and minimum length of retained sequences=40. Final total sequence yield per individual ranged from 10,665,214 to 39,607,344, with an average of 22,251,309 (Appendix S1). To create the cpDNA data set, reads from each individual (post-capture samples for Sets 1 and 2, pre- and post-capture samples for Set 3) were assembled in Geneious v. 9.0 to an annotated *Hordeum vulgare* chloroplast genome, Genbank accession NC_008590 (Saski et al., 2007). The post-capture vs. pre-capture cpDNA sequence profiles differed consistently, in that the coverage depth from the pre-capture libraries was far more even across the genome. The DNA reads that assembled to the chloroplast genome from each sample were saved (disassembled) in a separate file, and reassembled to the same *H. vulgare* chloroplast genome; this second assembly step reduced alignment artifacts involving the ends of reads in some regions. The 76 chloroplast genome assemblies were aligned using MAFFT in Geneious 10.0. Alignments were slightly manually edited, using the *H. vulgare* annotations as landmarks.

To assemble the nuclear-locus data sets, the 1478 *Hordeum vulgare* coding sequences used in the bait design pipeline were used as reference sequences for read-mapping using HybPiper (Johnson et al., 2016). The relative coverage of each locus across the taxa was visualized using the HybPiper R script gene_recovery_heatmap.R (Johnson et al., 2016) (Appendix S2). The individual exon-based locus data sets were aligned using MAFFT in Geneious 10.0. An initial rapid visual triage led to rejection of 263 loci with very poor alignments, extensive length variation, and/or any missing taxa. The remaining 1121 alignments were assessed by eye; minor ambiguities and misalignments were edited in Geneious 10.0, and complicated ambiguities were masked and excluded from phylogenetic analyses. These data sets were analyzed individually using RAxML version 8.2.10 (Stamatakis, 2014). The best tree for each locus, out of 20 runs under GTR+gamma model, was examined for anomalies—in particular, either very long branches, or splitting of taxa into multiple clades in a manner suggesting paralogy. These observations, in combination with output from HybPiper identifying potential paralogs, were used to eliminate an additional 75 loci, yielding a final total of 1046 data sets, all with 76 taxa.

Phylogenetic analyses of all cpDNA and nuclear-locus data sets, including tree searches and estimates of bootstrap support, were done with RAxML version 8.2.10 (Stamatakis, 2014). In general, searches were carried out under the GTR+gamma model of sequence evolution (Lanave et al., 1984; Yang, 1994), and bootstrap support was estimated using the fast-bootstrap option under the GTRCAT model (Stamatakis, 2006). Bootstrap tree files were used to generate 50% majority-rule consensus trees in RAxML (Aberer, Pattengale, and Stamatakis, 2010; Stamatakis, 2014). The full 1046-locus data set was also analyzed using a gene-tree approach with ASTRAL III (Zhang et al., 2018).

Three analyses of the chloroplast genome data were done: the entire data set including non-coding regions; only the protein-coding sequences, partitioned by gene; and only the protein-coding sequences, partitioned by codon position. Tree searches were done using a GTR+gamma model of evolution with the best tree saved from 50 independent runs, and bootstrap support was based on 100 replicates under the GTRCAT model.

The 1046 nuclear-locus data sets were analyzed individually using a GTR+gamma model of evolution, with the best tree saved from 50 independent runs for each locus (results not shown). Bootstrap analyses were carried out for each locus using the GTRCAT model (Stamatakis, 2006) with the number of replicates for each locus determined by the autoMRE stopping criterion (Pattengale et al., 2010).

The 1046 data sets were concatenated using SequenceMatrix (Vaidya, Lohman, and Meier, 2011). Tree searches were carried out using 50 independent runs under the GTR+gamma model, and bootstrap estimates were based on 250 replicates under the GTRCAT model. The analyses were first done on the unpartitioned data set, and then repeated with the data set partitioned by locus. In the partitioned analysis, the model was held constant across loci (GTR+gamma), but the model parameters were estimated independently for each locus. Phylogenetic relationships were also estimated using ASTRAL III (Zhang et al., 2018) with multilocus bootstrapping (Seo, Kishino, and Thorne, 2008). We ran an initial analysis using the best ML trees with weak branches (<10% bootstrap support) collapsed. Based on that tree, species with multiple representatives were identified as either a single lineage or as polyphyletic. The former were treated as a single terminal while the latter (*Eremopyrum distans*, *Hordeum stenostachys*, and *Secale strictum*) were treated as two species each, and the analyses were rerun.

The *Hordeum vulgare* sequences from each of the 1046 individual data sets were mapped to the barley genome online using BARLEYMAP (Cantalapiedra et al., 2015); 895 were unambiguously mapped to one contig associated with one of the seven barley chromosomes (Appendix S3, S4). These 895 loci were divided into seven subsets based on their *Hordeum* chromosome locations, and into 14 subsets based on chromosome-arm locations using centromere information from Mascher et al. (2017). While large portions of Poaceae and Triticeae genomes are broadly colinear (Devos and Gale, 2000), there have been chromosomal translocations during the history of the Triticeae (e.g., Devos et al., 1993; Martis et al., 2013; Danilova et al., 2017; Naranjo, 2019), so our chromosome/chromosome-arm based locus groups do not fully correspond to map locations throughout the tribe. However, the *Hordeum* maps do provide a reasonable heuristic approach to dividing loci into subsets to examine phylogenetic conflict. The 14 chromosome arm data sets were each analyzed using 50 independent searches under the GTR+gamma model, and bootstrap estimates were based on 500 replicates under the GTRCAT model. These results were compared to the concatenated data set of all 895 mapped loci, which was partitioned by locus and analyzed using 25 independent runs under the GTR+gamma model, with bootstrap estimates based on 250 replicates under the GTRCAT model.

Finally, to provide a more tractable visual comparison of the concatenated-data tree to the data-subset trees, we reanalyzed reduced-taxon data sets with a subset of 24 taxa selected to represent the diversity of the group (Appendix 1). First, a 24-taxon ML tree was obtained from a concatenation of all 895 *Hordeum*-mapped loci using a GTR+gamma model. Separate 24-taxon trees were then obtained for the seven chromosome subsets, for the 14 chromosome-arm subsets, and for a matched 24-taxon cpDNA data set. Bootstrap values were obtained using 250 replicates (895-locus data set) or 100 ML replicates (all data subsets and the cpDNA data set) using a GTRCAT model. The chromosome- and cpDNA-based bootstrap trees were compared to the 895-locus bootstrap tree, and conflicting relationships supported by at least 70% bootstrap support on at least one subset were shown as reticulations on the 895-locus tree.

## RESULTS

The chloroplast-genome phylogenetic tree (Fig. 1) is generally strongly supported. There are no notable differences between the trees based on the data set that includes non-coding regions (not shown) vs. those based on coding sequences only, whether partitioned by codon position (Fig. 1) or by gene (not shown). The tree based on the full data set has slightly higher bootstrap support. Differences among the three trees are limited to very short branches with low support, including (1) monophyly vs. paraphyly of *Taeniatherum caput-medusae* and *Aegilops speltoides* relative to the *Aegilops*/*Amblyopyrum*/*Triticum* clade; (2) the exact placement of *Dasypyrum villosum* within the *Dasypyrum*/*Pseudoroegneria*/*Thinopyrum* (Das/Pse/Thi) clade; (3) monophyly vs. paraphyly of *Agropyron cristatum* relative to *A. mongolicum*; and (4) the exact relationships among the four *Psathyrostachys juncea* accessions.

**Figure 1.**
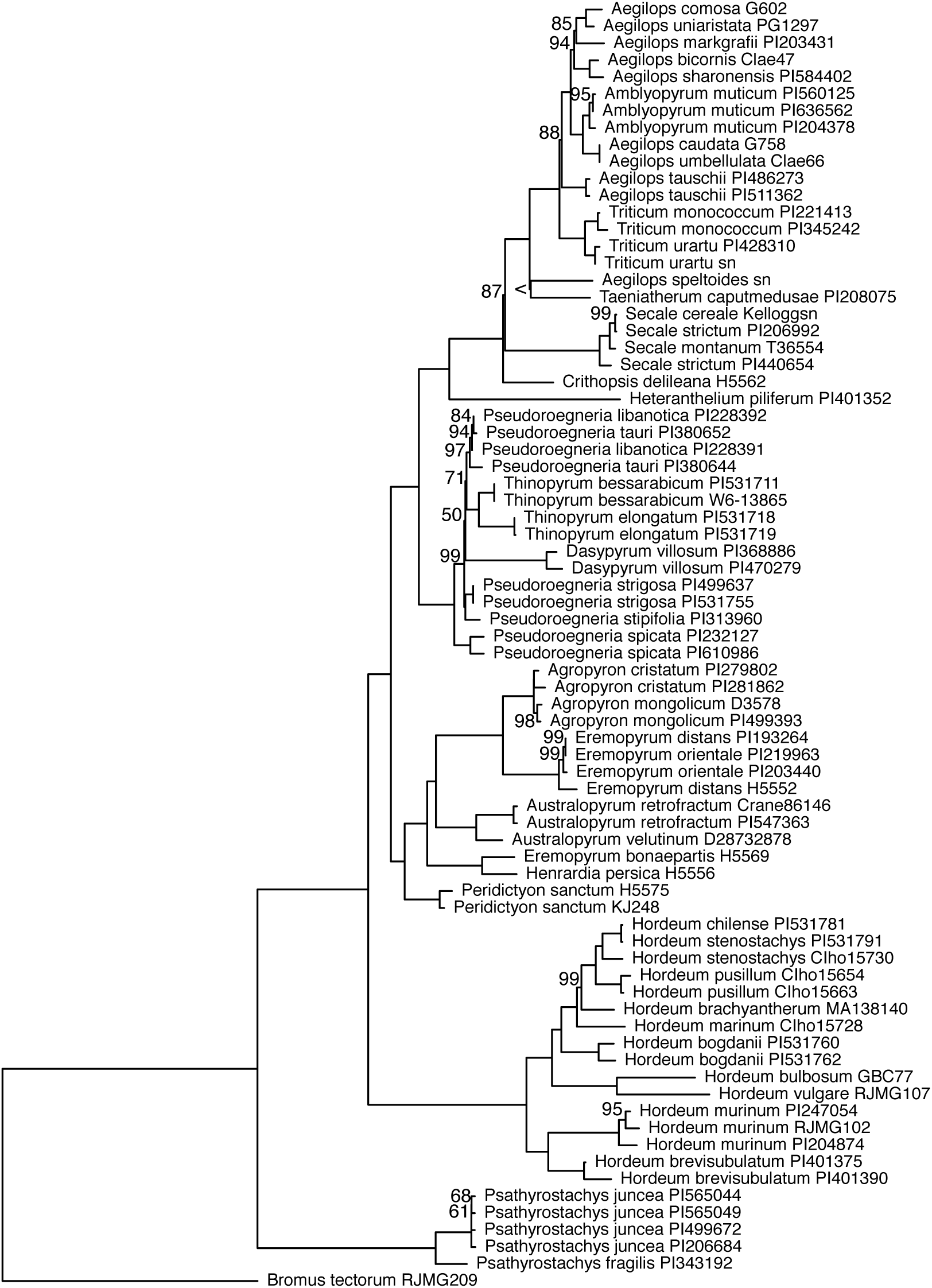
Phylogenetic estimate based on chloroplast genome coding sequences; this is the best of 50 trees from a ML search with a GTR+gamma model of evolution, partitioned by codon position. Bootstrap support was based on 100 ML replicates under the GTRCAT model. Unmarked nodes have 100% bootstrap support; numbers represent bootstrap values 50–99%; < represents bootstrap support under 50%.

While many of the individual nuclear-locus trees support the monophyly of the genomic genera, most show only weak support for deeper relationships among genera (results not shown; available in the online archive). Including introns in the alignments would yield more variation, likely leading to better resolution and stronger support, but introns can’t be reliably aligned across the entire tribe without extensive manual intervention. Earlier analyses of manually-aligned, cloned-sequence data sets (e.g., Mason-Gamer, Weil, and Kellogg, 1998; Mason-Gamer, 2001; Helfgott and Mason-Gamer, 2004) demonstrate phylogenetically useful variation in introns along with extensive length variation, sometimes involving insertions and deletions of transposons tens or hundreds of basepairs in length. Such frequent and substantial length changes confound automated alignment methods—even the exon-only data sets required visual inspection and numerous adjustments to accommodate indels.

The three trees based on 1046 nuclear loci—the concatenated analyses partitioned by locus (Fig. 2) and unpartitioned (not shown), and the ASTRAL III species tree (Appendix S5)— have much in common. The two ML trees are identical in topology. The partitioned analysis yields slightly higher bootstrap support; the proportions of nodes with 100% support in the partitioned vs. unpartitioned results are 67/73 vs. 60/73, respectively. The weakest relationship on both ML trees places *Henrardia persica* weakly (50%) at the base of a clade (hereafter referred to as the “wheats-rye” clade) that includes the wild wheats (*Triticum*, *Aegilops*, and *Amblyopyrum*), rye relatives (*Secale*), and three other genera (*Taeniatherum*, *Thinopyrum*, and *Crithopsis*). Though not evident from the ML topologies, there is underlying bootstrap support for an alternative placement of *H. persica* in a weak clade with *Agropyron* and *Eremopyrum* (Agr/Ere/Hen clade); the ASTRAL III tree (Appendix S5) strongly supports this relationship (posterior probability=1.0). The concatenated ML trees include a *Dasypyrum*, *Heteranthelium*, and *Australopyrum* group (100% in the partitioned analysis and 54% in the unpartitioned analysis), while the ASTRAL III tree instead places *Heteranthelium* and *Dasypyrum* at the base of the “wheats-rye” clade (1.0 posterior probability). Overall, most of the nuclear locus-based relationships are concordant across the three analyses, although a few weak support values and/or differences between the ML and ASTRAL III trees point to instability of the placement of *Dasypyrum*, *Heteranthelium*, and *Henrardia*, and the identity of the closest relatives to the “wheats-rye” clade.

**Figure 2.**
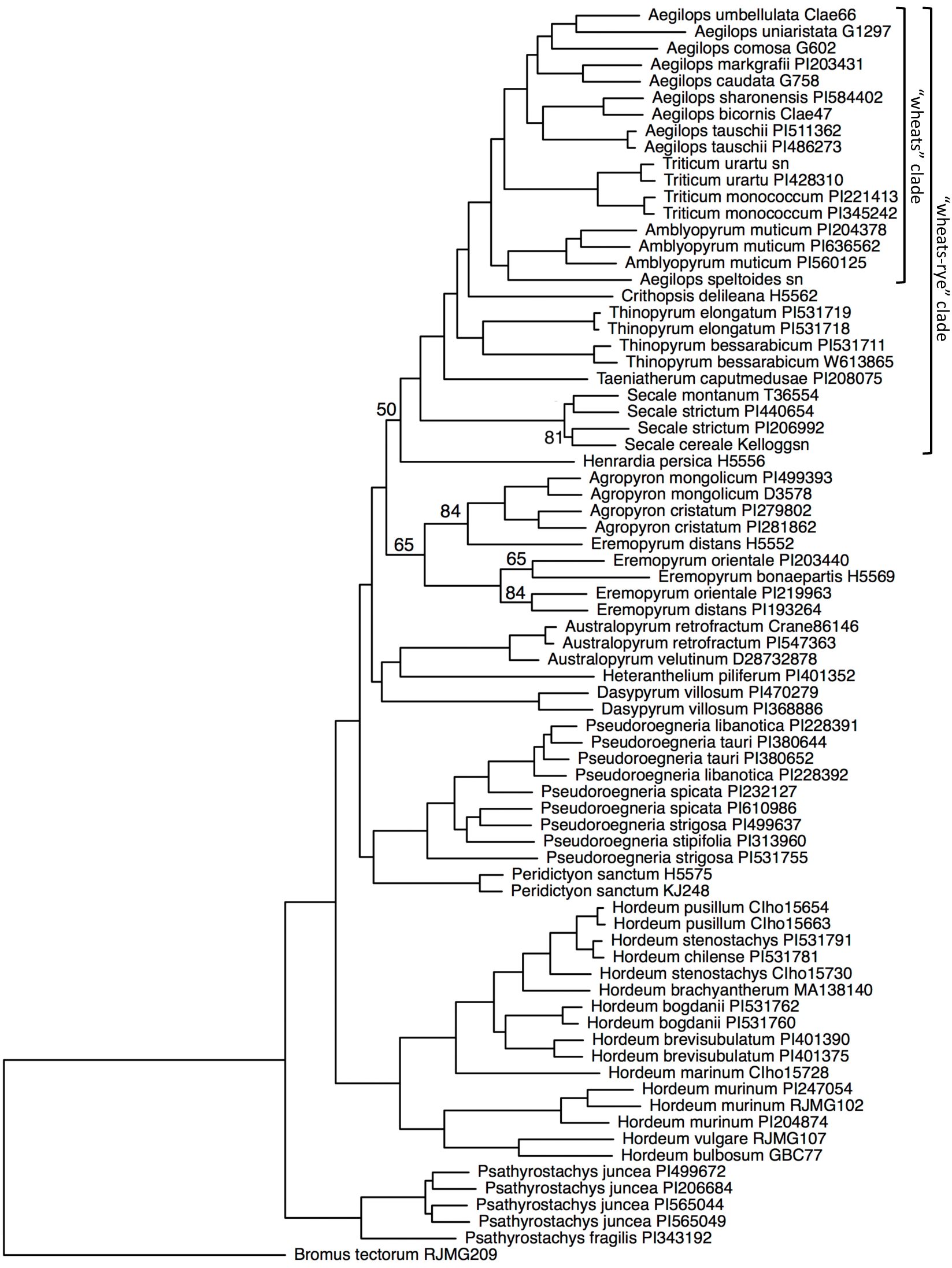
Phylogenetic estimate based on 1046 concatenated nuclear loci; this is the best of 50 trees from a ML search with a GTR+gamma model of evolution, partitioned by locus. Bootstrap support was based on 250 ML replicates under the GTRCAT model. Unmarked nodes have 100% bootstrap support; numbers represent bootstrap values 50–99%. The two labeled clades are referred to in the text.

Comparisons between the chloroplast-genome (Fig. 1) and nuclear-locus (Fig. 2, Appendix S5) trees reveal a few similarities and multiple major differences. Well-supported similarities include: (1) the basal position of *Psathyrostachys*, followed by *Hordeum*, relative to the designated outgroup *Bromus tectorum*; (2) the close relationship of *Crithopsis*, *Taeniatherum*, and *Secale* to the “wheats”; and (3) a close relationship between *Agropyron* and *Eremopyrum*. Reciprocally well-supported differences include, for example: (1) *Dasypyrum*, *Pseudoroegneria*, and *Thinopyrum* form a clade on the cpDNA tree, but are widely polyphyletic on the nuclear-locus trees; (2) *Agropyron*, *Australopyrum*, and *Eremopyrum* form another cpDNA clade, whereas *Australopyrum* is separated from the *Agropyron*/*Eremopyrum* clade on the nuclear-locus trees; and (3) the relative placements of *Crithopsis*, *Taeniatherum*, and *Secale* differ with respect to the base of the “wheats” clade. A tree-to-tree comparison generated using the cophylo function in the R package phytools (Revell, 2012) provides a visual comparison of the cpDNA vs. the concatenated partitioned ML tree (Appendix S6).

Analyses of subsets of loci, grouped according to their map location on *Hordeum* chromosomes (results not shown) and chromosome arms (Appendix S7), further highlight the aforementioned uncertainty of nuclear-locus relationships involving *Dasypyrum*, *Heteranthelium*, and *Henrardia*. Comparisons among the data subset trees point to additional uncertainties as well—for example, the positions of *Peridictyon* and *Crithopsis* are congruent with 100% support on the full concatenated and ASTRAL III trees, but some of the subset trees show strong conflict with respect to these two taxa. Figure 3 summarizes these and other selected differences among 17 trees—concatenated nuclear, ASTRAL III, chromosome arm subsets, and cpDNA.

**Figure 3.**
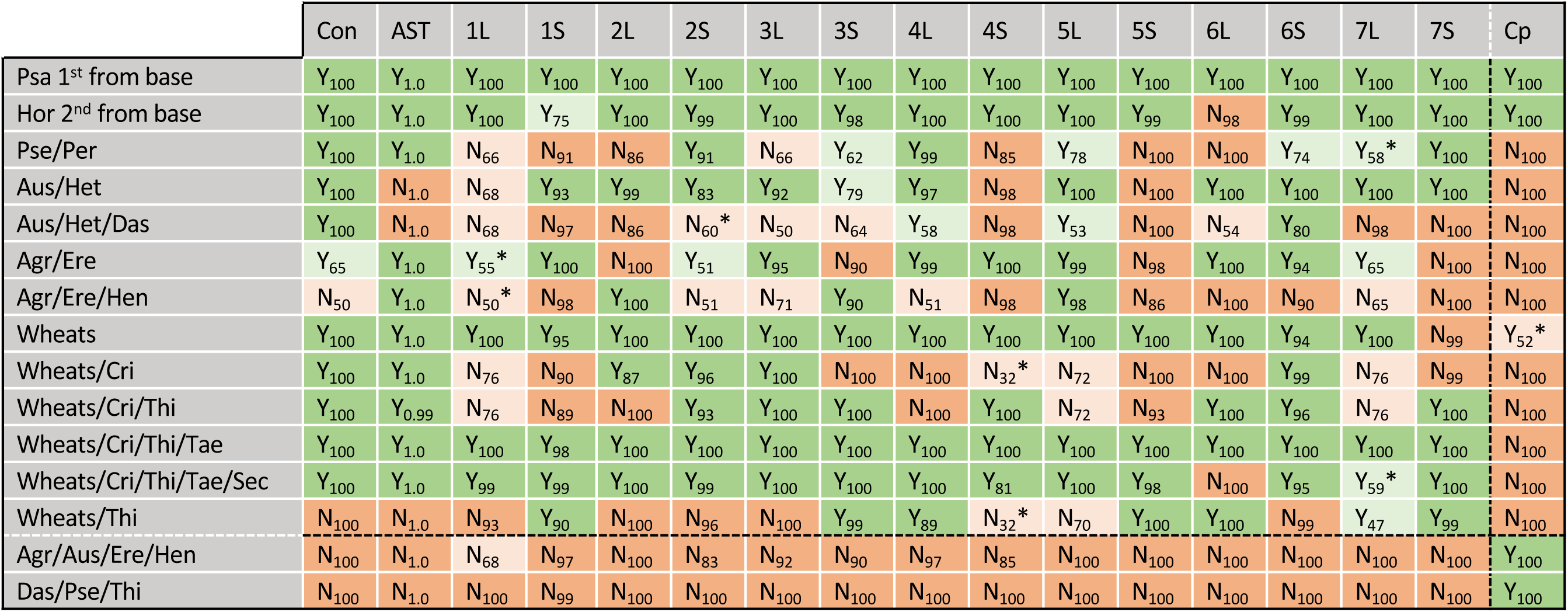
Summary of consensus and conflict among trees (top row) for selected nodes (le? column). Nodes present on a given ML tree are indicated by green cells with “Y” and the ML bootstrap support for that rela$onship; posterior probabili$es (pp) are provided for the ASTRAL III tree. Nodes that are not present are indicated by orange cells with “N” and support for the strongest conflic$ng node. Asterisks mark cases where the highest bootstrap support is not consistent with the best ML tree topology. Darker shades = bootstrap/pp support above 80%/0.8. DoSed lines highlight two unique cpDNA nodes (last two rows) and the cpDNA tree (right column). Tree abbrevia$ons: Con = concatenated nuclear loci, par$$oned by locus; AST = ASTRAL tree from nuclear loci (Appendix S5); 1–7L and 1–7S = concatenated loci from, respec$vely, the long and short arms of chromosomes 1–7 (Appendix S7); Cp = cpDNA coding-sequences, par$$oned by codon posi$on (Fig. 1). Node abbrevia$ons: Psa = *Psathyrostachys*; Hor = *Hordeum*; Pse = *Pseudoroegneria*; Per = *Peridictyon*; Aus = *Australopyrum*; Het = *Heteranthelium*; Das = *Dasypyrum*; Agr = *Agropyron*; Ere = *Eremopyrum*; Hen = *Henrardia*; Wheats = *Tri8cum*, *Aegilops*, and *Amblyopyrum*; Cri = *Crithopsis*; Thi = *Thinopyrum*; Tae = *Taeniatherum*; Sec = *Secale*.

While the 14 chromosome-arm trees differ from the concatenated and/or ASTRAL III tree in various ways (Fig. 3), the tree representing the long arm of chromosome 6 (6-long; Appendix S7, tree k) stands out in several respects. First, it is the only nuclear-locus tree that does not group *Secale* as part of the “wheats-rye” clade defined earlier (see row “Wheats/Cri/Thi/Tae/Sec” in Fig. 3). Instead, the 6-long tree is unique in placing *Secale* in a well-supported (100% bootstrap) clade with *Agropyron* and *Eremopyrum*. While the 6-long tree agrees with all other trees in its support of *Psathyrostachys*’s basal position, it is unique in that it does not identify *Hordeum* as the next-diverging lineage. Instead, the branching order on the 6-long tree is *Psathyrostachys*, *Peridictyon*, the unique *Secale*/*Agropyron*/*Eremopyrum* clade mentioned above, and then *Hordeum*.

The reduced-taxon results (Fig. 4) allow for a more simplified graphical depiction of the ways in which the cpDNA tree and the chromosome-based trees differ from the genome-wide data set tree (and, indirectly, from one another). Like the all-taxon analyses, this depiction highlights a few stable relationships, e.g., the basal position of *Psathyrostachys* relative to the designated outgroup *Bromus* on all trees. The placement of *Hordeum* inside of *Psathyrostachys* is supported by all but the chromosome 6 and 6-long trees, and the “wheats-rye” clade mentioned above is supported by all but the chromosome 6/6-long and cpDNA trees. On the other hand, the primary collective visual result is one of differences among trees, involving most of the other nodes. A particularly striking example involves the closest relative of the “wheats-rye” clade—the dashed lines G, H, O, P, and Q (Fig. 4) show the taxa or clades that occupy this position, with moderate to strong bootstrap support, on at least one chromosome-based tree.

**Figure 4.**
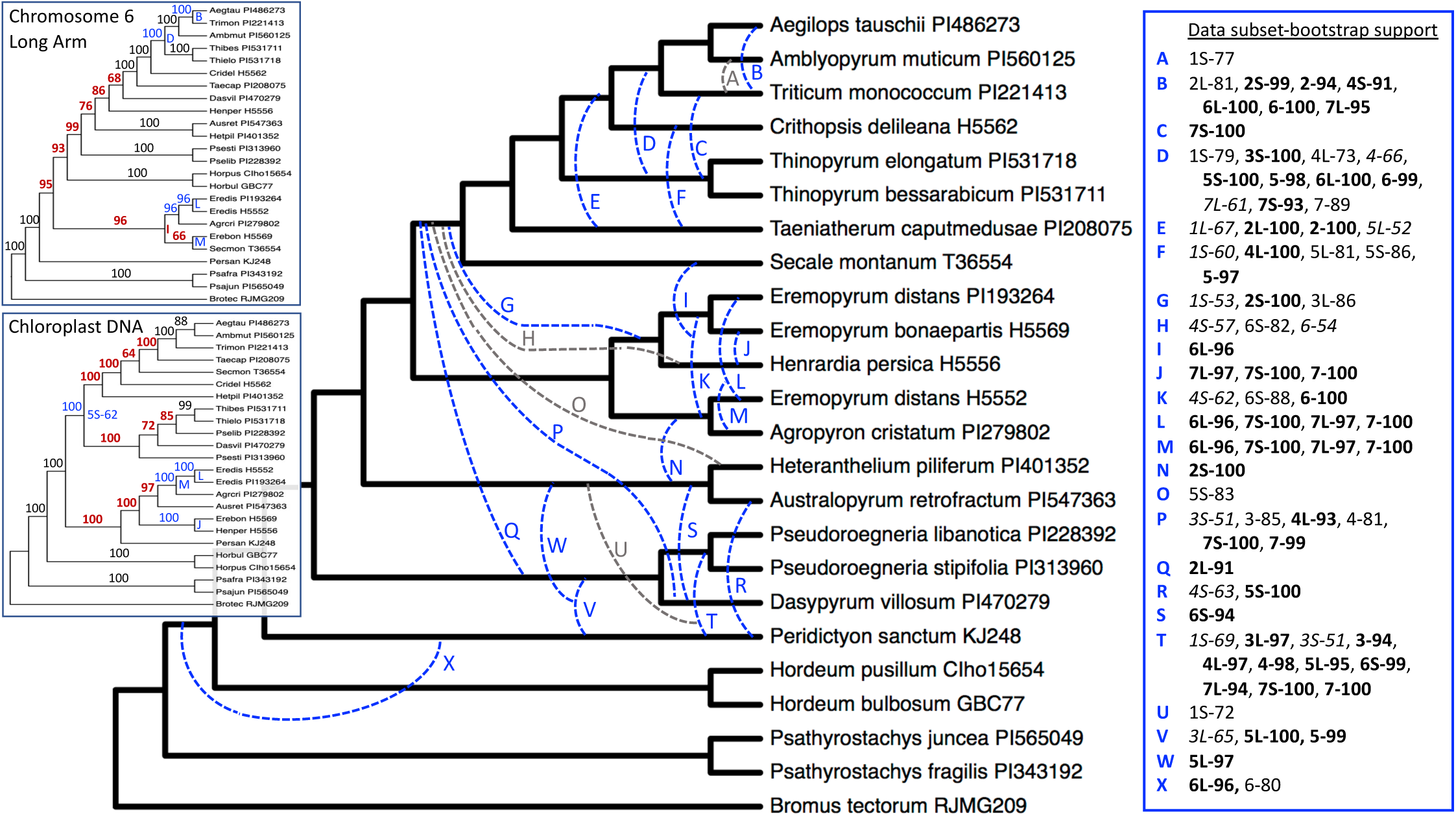
Chromosome-based trees vs. the concatenated nuclear-locus tree for 24 taxa; a few multistep conflicts could not be depicted in this format. Main tree: concatenation of 895 loci mapped to the barley genome, the best of 100 ML searches with a GTR+gamma model, partitioned by locus. All nodes have 100% bootstrap support based on 250 ML replicates under the GTRCAT model. Dashed lines: alternative relationships appearing on at least one chromosome-based dataset with ≥ 70% support based on 100 ML bootstrap replicates under the GTRCAT model (blue >90%; gray 70–89%). Uppercase letters refer to the table, which lists the chromosome-based trees on which that relationship appears, and its support on that tree (boldface >90%; plain text 71–89%; italic 50–70%). Table abbreviations: 1–7, 1L–7L, and 1S– 7S = concatenated loci representing, respectively, *Hordeum* chromosomes 1–7 and the long and short arms of chromosomes 1–7. Insets: the 6L tree and the cpDNA tree have multiple relationships that can’t be easily depicted on the main tree. Inset bootstrap values: black = concordant with the main tree; blue = conflict with the main tree but concordant with one or more other chromosome subsets; red = unique to that tree.

## DISCUSSION

The present phylogenetic analysis of a large multilocus nuclear data set builds upon a long history of molecular phylogenetic studies of the Triticeae. Many of the earlier analyses were based on individual loci, and focused on conflicting phylogenetic signal among gene trees (Kellogg, Appels, and Mason-Gamer, 1996; Mason-Gamer and Kellogg, 1996a; Mason-Gamer, 2005). Others discussed phylogenetic estimates from combined multilocus data sets, while still addressing apparent conflicting signal among them (Seberg and Petersen, 2007; Escobar et al., 2011). We take the latter approach on a larger scale through analyses of, and comparisons among, the cpDNA genome data set, the full nuclear-locus data set, and chromosome-based data subsets. We find that (1) the well-supported cpDNA phylogeny is in accord with previous cpDNA-based estimates; (2) most relationships are consistent and well-supported across multiple analyses of the full nuclear-locus data set, although (3) by the same criteria, some taxa remain poorly resolved by this data set; (4) although chloroplast and nuclear trees share a few points of agreement, they are largely in conflict; and (5) comparisons among chromosome arm-based trees further highlight a combination of stable, unstable, and incongruent relationships, and reveal that a portion of chromosome 6 is especially incongruent with the rest.

Recovering a chloroplast genome-based phylogenetic tree is straightforward, in principle. The largely uniparental, clonal inheritance of the haploid chloroplast genome, and its overall structural stability, minimize complications that might mislead phylogenetic analyses. The main problem is typically a lack of phylogenetic resolution and/or low support, which can be addressed with the increasing ease of obtaining large, multi-gene alignments. The present cpDNA analysis provides a matched-sample comparison with the multilocus nuclear phylogeny. It builds directly upon previous analyses of a comparable taxon sample but with far less data (e.g., Mason-Gamer, Orme, and Anderson, 2002), and it complements a more recent chloroplast-genome analysis involving a different set of individuals and extensive intraspecific sampling (Bernhardt et al., 2017). The present cpDNA tree generally confirms the relationships uncovered by Bernhardt et al. (2017), and includes accessions of two additional, monotypic genera: *Crithopsis delileana*, which is grouped with the *Secale*/*Taeniatherum*/”wheats” clade; and *Peridictyon sanctum*, grouped with the *Agropyron*/*Eremopyrum*/*Australopyrum*/*Henrardia* clade. At this point, we conclude that the cpDNA-based relationships among the Triticeae genera are largely settled. However, an interesting result from Bernhardt et al. (2017), which we are unable to address with our present sampling, is that *Psathyrostachys* fell outside of Triticeae + *Bromus* (presumed closest outgroup), a result consistent with some broader cpDNA-based analyses of the subfamily (e.g., Catalán, Kellogg, and Olmstead, 1997; Orton et al., 2021). This would suggest chloroplast exchange early in the evolution of the clade, wherein *Psathyrostachys* captured a Bromus chloroplast genome through introgression (as favored by Bernhardt et al., 2017), or the reverse.

In contrast to the relative ease of analyzing cpDNA data sets, phylogenetic analyses of multilocus nuclear data sets are impacted by complications inherent in biparentally-inherited, recombining genomes. Recombination among loci, for example, can lead to differences among nuclear gene trees when combined with incomplete lineage sorting and/or introgression. Quantifying differences among individual locus trees can potentially shed light on these processes, but few of our individual locus trees provided enough well-supported resolution to allow detailed comparisons among them; thus, we looked for phylogenetic inconsistencies by comparing the concatenated ML and ASTRAL III gene-tree analyses of the all-locus data sets to one another, and to individual analyses of chromosome-based data subsets.

Analyses of nuclear loci can be further complicated by polyploidy and gene duplication, and the potential assumption of orthology when homeologs or paralogs are actually being compared. In this study, homeology is not expected to be an issue because, while polyploidy is widespread throughout the Triticeae, these analyses include only diploid (2n=14) representatives. Complications due to paralogy are far more difficult to rule out; some of our single-locus data sets could include paralogs. The bait-design pipeline (Weitemier et al., 2014) is devised to minimize targeting genes with multiple paralogous copies, and individual gene trees were subsequently assessed visually for splits suggestive of paralogy, and analytically using HybPiper paralog warnings (Johnson et al., 2016). With this, we hope to have minimized the effects of paralogous gene copies on the phylogenetic results, but given a history of gene duplication events within diploid Triticeae genomes (e.g., Wang et al., 2022), we can’t entirely rule them out.

The Triticeae taxonomic system we follow, as discussed in the sample description, is based on the genomic taxonomic system of Löve (1984), which defines genera based on chromosome pairing. DNA sequence-based phylogenies to date, including those presented here, are generally consistent with the genomic system; in other words, on our cpDNA and nuclear DNA trees, most of the named genera form monophyletic groups. Two exceptions, *Eremopyrum* and the genera within the “wheats” clade, are discussed below. While the genomic system of classification defines generic *boundaries*, it does not address relationships *among* genera. There have been attempts to apply higher-lever intratribal taxonomic subdivisions (e.g., reviewed in West, McIntyre, and Appels, 1988), but they are not widely applied. While most of the intergeneric relationships based on the full nuclear-locus data sets presented here are robust to different analysis methods (i.e., they were well-supported in consistent relationships on the partitioned and unpartitioned ML trees, and on the ASTRAL III species tree), the relationships on the cpDNA tree broadly conflict with those on the nuclear trees. Thus, these results don’t especially enlighten attempts to define taxonomic subtribes within the Triticeae, although the one major feature shared by the cpDNA and nuclear trees— the paraphyletic *Psathyrostachys*/*Hordeum* assemblage vs. the remainder of the genera—does roughly correspond with, respectively, subtribes Hordeinae and Triticinae as recently summarized and defined by Feldman and Levi (2023a). Ultimately, however, any proposed subdivisions in the tribe are complicated not only by remaining phylogenetic uncertainties among genera, but also because subtribe divisions don’t easily accommodate extensive reticulation events—hybridization and allopolyploidy—that span the tribe and blur boundaries among proposed subgroups.

Only a few taxa were unstable across the three all-locus nuclear analyses, being weakly-supported on one or more trees, and/or variable among the three trees. One example is *Henrardia*, a lesser-known genus of two annual species, native to western and central Asia (Hubbard, 1946). It, along with several other small Triticeae taxa, is listed among the poorly-understood “orphan genera” of Feldman and Levy (2023b), and in our analyses at least, it remains ambiguous. Its unique morphology has led to divergent views about *Henrardia*’s appropriate placement within the tribe (Frederiksen, 1993), ranging from classification in its own separate subtribe (Löve, 1984) to placement within *Aegilops* (Kellogg, 1989). In earlier gene-tree analyses, *Henrardia*’s position was generally poorly supported (e.g., Hsiao et al., 1995; Mason-Gamer, 2001, 2005; Seberg and Petersen, 2007), although one analysis suggested a unique relationship to *Secale* (Fan et al., 2013). A concatenated analysis of 27 loci (Escobar et al., 2011) placed *Henrardia* with *Agropyron* and *Eremopyrum* (Agr/Ere/Hen clade), although that group was recovered on only one of their 27 individual gene trees. In agreement with Escobar et al. (2011), our ASTRAL III tree recovered the same Agr/Ere/Hen clade with strong support (99%), but this relationship is only weakly supported by the ML unpartitioned analysis, and is weakly contradicted by the ML partitioned analysis, which instead places *Henrardia* at the base of the previously mentioned “wheats-rye” clade. Moreover, there is considerable ambiguity among the 14 chromosome-arm based data subsets (see Fig. 3, row “Agr/Ere/Hen”). Only three support the Agr/Ere/Hen clade, albeit with 90–100% ML bootstrap support (Appendix 7, trees c, f, and i). Among the eleven that do not, eight place *Henrardia* in the same position as the ML partitioned analysis (basal to the ”wheats-rye” clade) with support ranging from 50% to 98% (Appendix 7, trees a, b, d, e, g, h, l, and m).

Potentially more misleading are relationships that are consistent and well-supported across all three analyses of the full nuclear-locus trees, with underlying instability only revealed by data subsets. Large, concatenated phylogenomic data sets can yield very high bootstrap values for incorrect relationships (e.g., Huang et al., 2021), including those that conflict among data subsets (e.g., Chan et al., 2020). Here, *Peridictyon* illustrates the latter phenomenon. It is another small, rather obscure monotypic genus (Seberg et al., 1991). As with *Henrardia*, earlier nuclear-locus analyses (e.g., Hsiao et al., 1995; Mason-Gamer, 2001; Helfgott and Mason-Gamer, 2004; Mason-Gamer, 2005; Seberg and Petersen, 2007; Fan et al., 2013) shed little light on *Peridictyon*’s placement—its position on individual gene trees varied widely, and support was low in most cases. Our analysis brings much more data to bear, and *Peridictyon* and *Pseudoroegneria* form a well-supported clade on all three full-data set trees. However, only seven of the fourteen chromosome-arm based trees recover this relationship (Fig. 3, row Pse/Per; Appendix S7, trees d, f, g, i, l, m, and m). Among the seven dissenting trees, *Peridictyon* is placed in various relationships with a range of support: e.g., it forms a clade with *Australopyrum* (100% support) on the chromosome 5 short-arm (5-short) tree Appendix 7, tree j), and it is near the base of the 6-long tree, just inside of *Psathyrostachys* (98%; Appendix 7, tree k). The 6-long data set itself comprises mixed information about the placement of *Peridictyon—*when this data set is further divided into four subsets (Appendix S8, S9), one subset reflects the near-basal position of *Peridictyon* with 100% support; one weakly reflects the *Peridictyon*/*Pseudoroegneria* relationship seen on the all-data tree and on half of the chromosome-arm trees; one uniquely places *Peridictyon* within an also-unique *Agropyron*/*Eremopyrum*/*Secale* clade with 100% support; and one uniquely (but very weakly) places *Peridictyon* with *Secale*. Taken together, the data indicate that there is no single phylogenetic placement that fully describes *Peridictyon*’s relationship to the rest of the tribe, even though there is dispersed, genome-wide phylogenetic signal placing it with *Pseudoroegneria*.

While we primarily focus on deeper phylogenetic relationships and conflict among genera, incongruence is also evident toward the tips of the tree (Figs. 5, 6). One case involves *Eremopyrum*, a small group of perennial species that includes both diploids and tetraploids (Frederiksen, 1991). Although *Eremopyrum* is not a focus of this study, even our small sample of diploids indicates considerable phylogenetic conflict (Fig. 5). While a polyphyletic *Eremopyrum* has been suggested by earlier analyses of individual nuclear gene trees, including *DMC1* (Petersen and Seberg, 2002), *Acc1* (Fan et al., 2013), beta-amylase (Mason-Gamer, 2005), a 27-locus data set (Escobar et al., 2011), and multiple cpDNA data sets (e.g., Mason-Gamer, Orme, and Anderson, 2002; Bernhardt et al., 2017), the present analysis highlights not only polyphyly, but a series of well-supported, conflicting relationships involving various groupings with *Henrardia* and *Agropyron* (Fig. 5). Both of these genera (among others) have been proposed as close relatives of *Eremopyrum*: e.g., *Agropyron* is morphologically similar overall to *Eremopyrum* (Clayton and Renvoize, 1986), and *Henrardia* and *Eremopyrum* share similar telocentric chromosomes, which are otherwise unusual in the tribe (Frederiksen, 1991). Our results simultaneously support both of these hypotheses, suggesting a hybrid origin of *Eremopyrum*. A larger sample across all three genera, with a particular focus on *Eremopyrum* tetraploids, would likely provide a clearer picture of their intertwined history. More generally, however this example further illustrates the complexity of even such a basic concept as relatedness in a widely reticulate group.

**Figure 5.**
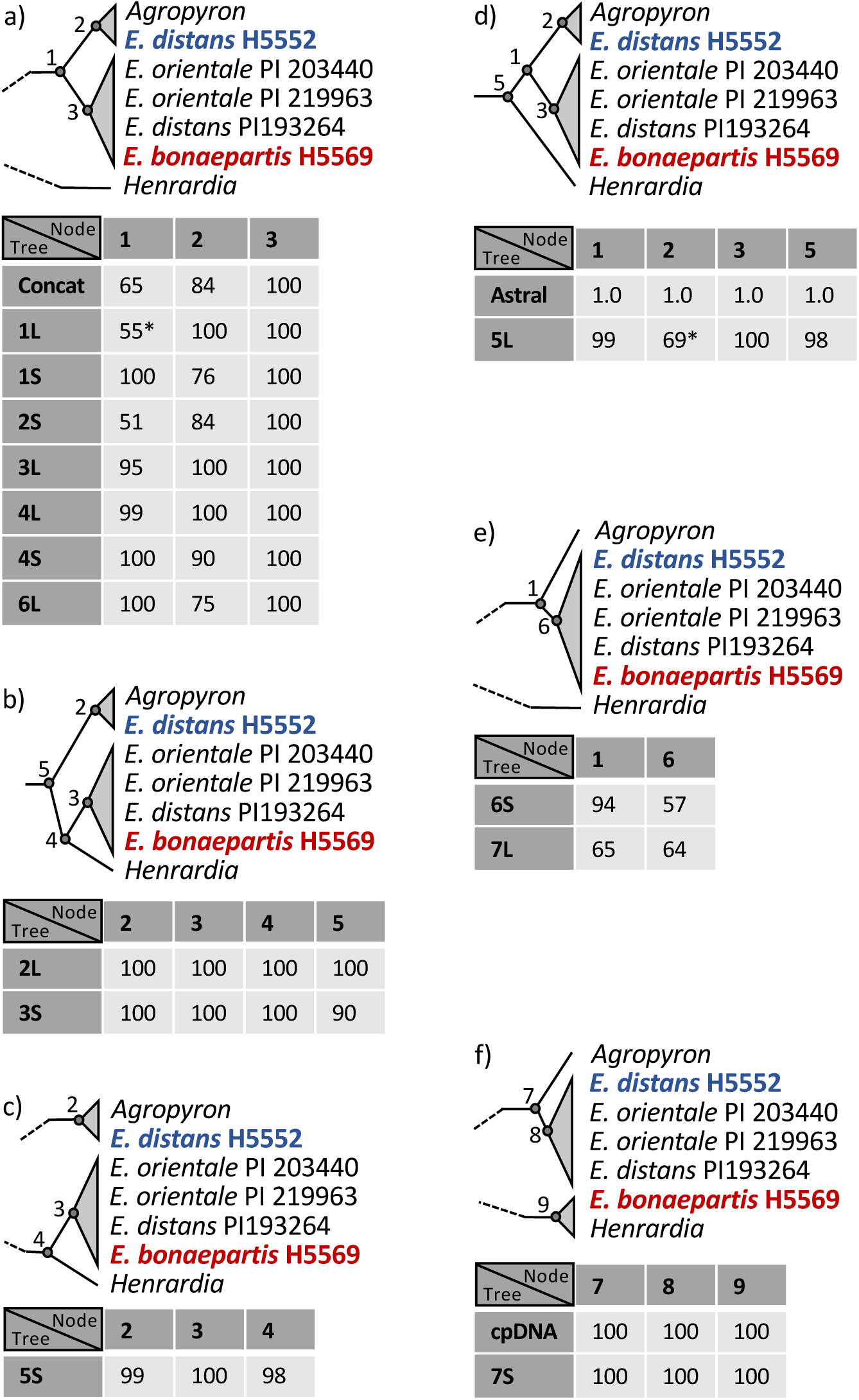
Summary of conflict and consensus among trees with respect to *Eremopyrum*, *Agropyron*, and *Henrardia*. Node numbers that appear on multiple panels (1–5) represent the same clade in each panel. The tables show ML bootstrap support for each node on each ML tree; posterior probability is shown for the ASTRAL III tree. Asterisks mark cases where the highest bootstrap support is not consistent with the best ML tree topologies. Tree abbreviations: Concat = concatenated nuclear loci (Fig. 2); Astral = ASTRAL III tree from nuclear loci (Appendix S5); 1–7L and 1–7S = concatenated nuclear loci representing, respectively, the long and short arms of *Hordeum* chromosomes 1–7 (Appendix S7); cpDNA = cpDNA tree (Fig. 1).

Another case of genus-level conflict involves the diploid wild wheat species (*Triticum*, *Aegilops*, and *Amblyopyrum*). This group has been examined at length using one or a few nuclear loci (Petersen et al., 2006; Golovnina et al., 2007), genome-wide SNPs (Edet et al., 2018), cpDNA genomes (Li et al., 2015), multilocus genomic data sets (Marcussen et al., 2014; Huynh et al., 2019) and genome-wide RNA sequencing (Glémin et al., 2019; Tanaka et al., 2020). Some of these studies specifically highlight the effects of past hybridization deep within the clade, but the number of events, and the hypothesized participants, differ among analyses. The conclusions of two of the more comprehensive analyses (Glémin et al., 2019; Huynh et al., 2019) differ in terms of the number of inferred hybridization events, but both highlight four ancient lineages, leading to *Amblyopyrum muticum* (= *Aegilops mutica*), *Ae. speltoides*, *Triticum*, and the remaining *Aegilops* species. Both propose that the multi-species *Aegilops* lineage arose through an ancient hybridization event involving *Triticum*, with either *Am. muticum* (Glémin et al., 2019) or *Ae. speltoides* (Huynh et al., 2019). They reach different conclusions about whether this event is largely sufficient to explain present reticulate patterns (Huynh et al., 2019) or whether multiple additional hybridization events were involved (Glémin et al., 2019). In any case, the variable relationships that we observe in our “wheats” clade (Fig. 6) involve the relative placements of the same four lineages—*Am. muticum*, *Ae. speltoides*, *Triticum*, and the rest of the *Aegilops* species—and are thus consistent with a history of ancient hybridization among the diploid species.

**Figure 6.**
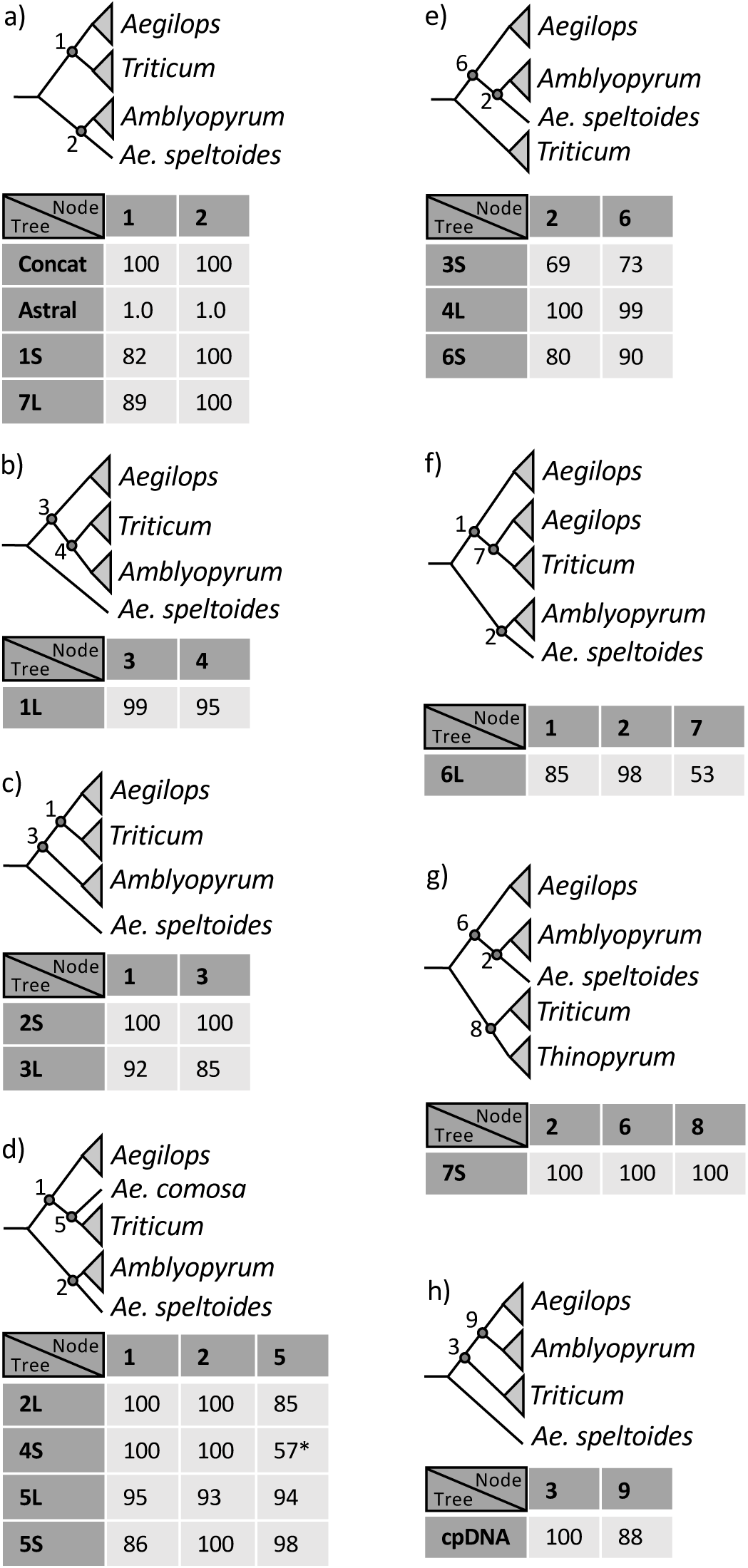
Summary of conflict and consensus among trees with respect to the “wheats” clade (*Aegilops*, *Triticum*, and *Amblyopyrum*). Node numbers that appear on multiple panels (1–3, 6) represent the same clade in each panel. The tables show ML bootstrap support for each node on each ML tree; posterior probability is shown for the ASTRAL III tree. An asterisk marks a case where the highest bootstrap support is not consistent with the best ML tree topology. Tree abbreviations: Concat = concatenated nuclear loci (Fig. 2); Astral = ASTRAL III tree from nuclear loci (Appendix S5); 1–7L and 1–7S = concatenated nuclear loci representing, respectively, the long and short arms of *Hordeum* chromosomes 1–7 (Appendix S7); cpDNA = cpDNA tree (Fig. 1).

The closest relative to the “wheats” clade is another point of conflict among nuclear data subsets. The full data trees—partitioned concatenated, unpartitioned concatenated, and ASTRAL III—all support *Crithopsis* as sister to the “wheats” clade (support = 100%, 74%, and 100%, respectively). However, only four of the 14 chromosome-arm trees support this relationship (support % = 87, 96, 100, 99; Appendix 7, trees c, d, e, and l), while six instead place *Thinopyrum* as sister to the “wheats” (support % = 90, 99, 89, 100, 100, 47; Appendix 7, trees b, f, g, j, k, m). The cpDNA tree, in contrast, places *Taeniatherum* as the closest relative to the “wheats,” just within the clade (support = 100%), followed by *Secale* (87%) and *Crithopsis* (100%). These results add to disparate relationships recovered in earlier studies based on nuclear loci, including: *Taeniatherum* at the base of the wheat clade, and *Thinopyrum* nested within it (Fan et al., 2013); *Thinopyrum* and *Crithopsis* nested within the wheat clade (Seberg and Petersen, 2007); a triphyletic wheat group with *Crithopsis* and *Thinopyrum* sister to each of two clades (Petersen et al., 2006; Mason-Gamer, 2013) and *Taeniatherum* nested within one of those (Mason-Gamer, 2013).

Studies of Triticeae systematics over decades broadly exemplify the history of the application of phylogenetic data in plants. The tribe or its members have been analyzed at length using data from morphological characters, life-history traits, interspecific crossability, isozyme mobility, chromosome pairing, genomic in situ hybridization, cpDNA restriction site and sequence variation, and DNA sequence variation from single-copy through highly-repetitive nuclear loci. Of these approaches, perhaps the most widely influential to our perception of the tribe have been the cytogenetic analyses of chromosome pairing, which have been used to circumscribe groups based on patterns of meiotic chromosome pairing and, in turn, have been instrumental in defining and naming Triticeae genera. These data, however, cannot resolve phylogenetic relationships among those genera, since lack of pairing does not contain hierarchical information. Molecular phylogenetic analyses were expected to clarify these deeper relationships, and indeed some have been reasonably well-settled—the early divergence of the *Pseudoroegneria* and *Hordeum* lineages, for example. However, after 25 years and many phylogenic analyses, we still find ourselves in much the same place; genomic genera are generally supported by molecular phylogenetic data, but many of the relationships among them remain unclear, due to a combination of conflict among data sets and weak or unstable clades. The increase in the amount of phylogenetic data applied to this question exacerbates both of these, at once increasing the support for incongruence among trees, while simultaneously failing to resolve certain problematic relationships. We close with a relevant quote from R. G. Olmstead and A. M. Bedoya (2019), who consider the impact of whole chloroplast genomes and extensive nuclear-genome sampling on the current state of plant phylogenetics: “*Where do we stand today? Whole genome sequencing, the ‘holy grail’ of molecular phylogenetics, provides us with more and powerful insights into plant phylogeny, but still leaves us pondering some of the same thorny questions we’ve struggled with for decades.*”

## Supporting information

Supplement S1-Samples

Supplement S2-Capture Success

Supplement S3-Chromosome List

Supplement S4-Chromosome Diagram

Supplement S5-ASTRAL Tree

Supplement S6-Tanglegram

Supplement S7-Chromosome Trees

Supplement S8-Chromosome 6 Loci

Supplement S9-Chromosome 6 Trees

## ACKNOWLEDGMENTS

The authors thank Kevin Weitemier (Oregon State University) and Mark Fishbein (Oklahoma State University) for help with bait design; Stefan Green and Kevin Kunstman (Research Resources Center, University of Illinois at Chicago) and Kevin Feldheim (Pritzker Lab, Field Museum of Natural History) for valuable advice on data collection; and input from John Freudenstein and two anonymous reviewers on a previous version of the manuscript. The research was supported by NSF-DEB 1354975.

## AUTHOR CONTRIBUTIONS

RJMG conceived the study. RJMG and DMW collected and analyzed the data. RJMG wrote the manuscript with input from DMW.

## DATA AVAILABILITY STATEMENT

The raw short-read sequence data were deposited in the NCBI SRA data base (Bioproject Accession PRJNA755334). A Zenodo data repository (DOI 10.5281/zenodo.7612904) contains: lists of 58,566 baits used for targeted sequence capture and 18,048 blocks from which the baits were designed; cpDNA alignment with coding and non-coding regions; cpDNA alignment of protein-coding sequences only, with individual genes defined as character sets; a concatenated alignment of 1046 captured nuclear loci, with individual loci defined as character sets; descriptive information about the locus data sets; individual locus bootstrap tree files and images; two concatenated alignments of 895 loci mapped to the barley genome—one with all taxa and one with a subset of 24 taxa—with chromosomes and chromosome arms defined as character sets.

## ORCID

*Roberta J. Mason-Gamer:* https://orcid.org/0000-0001-9116-1640

*Dawson M. White:* https://orcid.org/0000-0002-0670-9390

## SUPPORTING INFORMATION

Additional supporting information may be found online in the Supporting Information section at the end of the article.

**Appendix S1.** Taxon names, accession numbers, library sets, library adaptors, barcodes, total number of reads, number of chloroplast DNA reads, chloroplast DNA shotgun reads (where applicable), and percent chloroplast genome coverage.

**Appendix S2.** Graphical representation of percent reference coverage for the 1478 *Hordeum* reference sequences (loci) that were used for read-mapping in HybPiper. Each column represents a locus. As explained in the text, y1046 of these were used in subsequent phylogenetic analyses. Each row is a sample, listed from top to bottom in the order shown in Appendix S1. The 21^st^ sample with noticeably poor coverage is an outgroup, *Avena fatua*, which was not included in the phylogenetic analyses.

**Appendix S3.** Chromosome map locations of the 895 loci that were mapped to a contig associated with one of the seven barley chromosomes, determined using BARLEYMAP (Cantalapiedra et. al. 2015).

**Appendix S4.** From Appendix S3, approximate positions within the barley genome of the 895 loci used in the concatenated chromosome and chromosome-arm (Appendix S7) analyses, determined using BARLEYMAP (Cantalapiedra et. al. 2015). Rounded rectangles represent chromosome arms; centromere location information is from Mascher et al. (2015).

**Appendix S5**. ASTRAL III species tree with 1046 ML gene trees with branches with less than 10% bootstrap support collapsed. Multiple individuals within species were assigned to single terminals, except three that appeared polyphyletic on the ASTRAL III lineage tree of all individuals (not shown): *Eremopyrum distans* (Eredis), *Hordeum stenostachys* (Horste), and *Secale strictum* (Secstr). Posterior probability values less than 1.0 are shown.

**Appendix S6.** Tanglegram comparison of the cpDNA tree (right) and the concatenated 1046-locus nuclear tree (left) generated using the cophylo function in the R package phytools; some of the differences in topology are discussed in the text.

**Appendix S7**, a–u. Individual trees from 21 subsets of loci corresponding to the 7 *Hordeum* chromosomes and 14 chromosome arms (Appendix S3, S4). Each tree represents the best ML tree out of 50 runs under a GTR+gamma model and partitioned by locus, with support from 500 bootstrap replicates under the GTRCAT model. The distinctive tree from the long arm of chromosome 6 (q) is discussed in more detail in the text.

**Appendix S8.** Loci mapped to the long arm of *Hordeum vulgare* chromosome 6 (Appendix S3), with closest-to-farthest from the *H. vulgare* centromere listed top-to-bottom. The 53 concatenated loci were analyzed together (Appendix S7, tree q) and separated into subsets a–d (Appendix S9).

**Appendix S9.** Maximum-likelihood analyses of four subsets of loci mapped to the long arm of *Hordeum* chromosome 6 (Appendix S8).

## Notes

https://zenodo.org/uploads/7612904

https://www.ncbi.nlm.nih.gov/sra/?term=PRJNA755334

### Competing Interest Statement

The authors have declared no competing interest.

## References

Aberer, A. J., N. D. Pattengale, and A. Stamatakis. 2010. Parallelized phylogenetic post-analysis on multi-core anchitectures. Journal of Computational Science 1: 107–114.

Asay, K. H., W. H. Horton, K. B. Jensen, and A. J. Palazzo. 2001. Merits of native and introduced Triticeae grasses on semiarid rangelands. Canadian Journal of Plant Science 81: 45–52.

Barkworth, M. E. 2000. Changing perceptions of the Triticeae. In S. W. L. Jacobs and J. Everett [eds.], Grasses: Systematics and Evolution, 110–120. Csiro, Melbourne.

Barkworth, M. E. 2005. Twenty-one years later: the impact of Löve and Dewey’s genomic classification proposal. Czech Journal of genetics and Plant Breeding 41 (special issue): 3–9.

Barkworth, M. E., and D. R. Dewey. 1985. Genomically based genera in the perennial Triticeae of North America: identification and membership. American Journal of Botany 72: 767– 776.

Barkworth, M. E., D. R. Cutler, J. S. Rollo, S. W. L. Jacobs, and A. Rashid. 2009. Morphological identification of genomic genera in the Triticeae. Breeding Science 59: 561–570.

Baum, B. R., J. R. Estes, and P. K. Gupta. 1987. Assessment of the genomic system of classification in the Triticeae. American Journal of Botany 74: 1388–1395.

Bernhardt, N. 2015. Taxonomic treatments of Triticeae and the wheat genus Triticum. In M. Molnár-Láng, C. Ceoloni, and J. Doležel [eds.], Alien Introgression in Wheat: Cytogenetics, Molecular Biology, and Genomics, 1–19. Springer International Publishing, Switzerland.

Bernhardt, N., J. Brassac, B. Kilian, and F. R. Blattner. 2017. Dated tribe-wide whole chloroplast phylogeny indicates recurrant hybridizations within Triticeae. BMC Evolutionary Biology 17: 141.

Cantalapiedra, C. P., R. Boudiar, A. M. Casas, E. Igartua, and B. Contreras-Moreira. 2015. BARLEYMAP: physical and genetic mapping of nucleotide sequences and annotation of surrounding loci in barley. Molecular Breeding 35: 13.

CataláN, P., E. A. Kellogg, and R. G. Olmstead. 1997. Phylogeny of Poaceae subfamily Pooideae based on chloroplast *ndh*F gene sequences. Molecular Phylogenetics and Evolution 8: 150–166.

Chan, K. O., C. R. Hutter, P. L. Wood, L. L. Grismer, and R. M. Brown. 2020. Larger, unfiltered datasets are more effective at resolving phylogenetic conflict: Introns, exons, and UCEs resolve ambiguities in Golden-backed frogs (Anura: Ranidae; genus *Hylarana*). Molecular Phylogenetics and Evolution 151: 106899.

Clayton, W. D., and S. A. Renvoize. 1986. Genera Graminum. Grasses of the World. Royal Botanical Gardens, London.

Danilova, T. V., A. R. Akhunova, E. D. Akhunov, B. Friebe, and B. S. Gill. 2017. Major structural genomic alterations can be associated with hybrid speciation in *Aegilops markgrafii* (Triticeae). The Plant Journal 92: 317–330.

Devos, K. M., and M. D. Gale. 2000. Genome relationships: the grass model in current research. The Plant Cell 12: 637–646.

Devos, K. M., M. D. Atkinson, C. N. Chinoy, H. A. Francis, R. L. Harcourt, R. M. D. Koebner, C. J. Liu, et al. 1993. Chromosomal rearrangements in the rye genome relative to that of wheat. Theoretical and Applied Genetics 85: 673–680.

Dewey, D. R. 1984. The genomic system of classification as a guide to intergeneric hybridization within the perennial Triticeae. *In* J. P. Gustafson [ed.], Gene manipulation in plant improvement, Proc. 16th Stadler genetics symposium, 209–279. Plenum Publishing Company, New York, NY.

Edet, O. U., Y. F. A. Gorafi, S. Nasuda, and H. Tsujimoto. 2018. DArTseq-based analysis of genomic relationships among species of tribe Triticeae. Scientific Reports 8: 16397.

Escobar, J. S., C. Scornavacca, A. Cenci, C. Guilhaumon, S. Santini, E. J. P. Douzery, V. Ranwez, et al. 2011. Multigenic phylogeny and analysis of tree incongruences in Triticeae (Poaceae). BMC Evolutionary Biology 11: 181.

Fan, X., L.-N. Sha, S.-B. Yu, D.-D. Wu, X.-H. Chen, X.-F. Zhuo, H.-Q. Zhang, et al. 2013. Phylogenetic reconstruction and diversification of the Triticeae (Poaceae) based on a single-copy nuclear *Acc1* and *Pgk1* gene. Biochemical Systematics and Ecology 50: 346–360.

Feldman, M., and A. A. Levy. 2023a. Taxonomy and evolution of the tribe Triticeae Dumort. *In* M. Feldman and A. A. Levy [eds.], Wheat Evolution and Domestication 10.1007/978-3-031-30175-9_2 Doi, 9–41. Springer International Publishing, Cham.

Feldman, M., and A. A. Levy. 2023b. Orphan genera of the subtribe Triticineae Simmonds. In M. Feldman and A. A. LEvy [eds.], Wheat Evolution and Domestication 10.1007/978-3-031-30175-9_5 Doi, 85–157. Springer International Publishing, Cham.

Frederiksen, S. 1991. Taxonomic studies in *Eremopyrum* (Poaceae). Nordic Journal of Botany 11: 271–285.

Frederiksen, S. 1993. Taxonomic studies in some annual genera of the Triticeae (Poaceae). Nordic Journal of Botany 13: 481–493.

Fu, L., B. Niu, Z. Zhu, S. Wu, and W. Li. 2012. CD-HIT: accelerated for clustering the next-generation sequencing data. Bioinformatics 28: 3150–3152.

GléMin, S., C. Scornavacca, J. Dainat, C. Burgarella, V. Viader, M. Ardisson, G. Sarah, et al. 2019. Pervasive hybridizations in the history of wheat relatives. Science Advances 5: eaav9188.

Golovnina, K. A., S. A. Glushkov, A. G. Blinov, V. I. Mayorov, L. R. Adkison, and N. P. Goncharov. 2007. Molecular phylogeny of the genus *Triticum* L. Plant Systematics and Evolution 264: 195–216.

Helfgott, D. M., and R. J. Mason-Gamer. 2004. The evolution of North American *Elymus* (Triticeae, Poaceae) allotetraploids: evidence from phosphoenolpyruvate carboxylase gene sequences. Systematic Botany 29: 850–861.

Hsiao, C., N. J. Chatterton, K. H. Asay, and K. B. Jensen. 1995. Phylogenetic relationships of the monogenomic species of the wheat tribe, Triticeae (Poaceae), inferred from nuclear rDNA (internal transcribed spacer) sequences. Genome 38: 211–223.

Huang, J., Y. Liu, T. Zhu, and Z. Yang. 2021. The asymptotic behavior of bootstrap support values in molecular phylogenetics. Systematic Biology 70: 774–785.

Hubbard, C. E. 1946. *Henrardia*, a new genus of the Gramineae. Blumea Suppl. 3: 10–21.

Huynh, S., T. Marcussen, F. Felber, and C. Parisod. 2019. Hybridization preceded radiation in diploid wheats. Molecular Phylogenetics and Evolution 139: 106554.

International Barley Genome Sequencing Consortium, T. 2012. A physical, genetic and functional sequence assembly of the barley genome. Nature 491: 711–717.

Johnson, M. G., E. M. Gardner, Y. Liu, R. Medina, B. Goffinet, A. J. Shaw, N. J. C. Zerega, and N. J. Wickett. 2016. HybPiper: extracting coding sequence and introns for phylogenetics from high-throughput sequencing reads using target enrichment. Applications in Plant Sciences 4: 1600016.

Kearse, M., R. Moir, A. Wilson, S. Stones-Havas, M. Cheung, S. Sturrock, S. Buxton, et al. 2012. Geneious Basic: an integrated and extendable desktop software platform for the organization and analysis of sequence data. Bioinformatics 28: 1647–1649.

Kellogg, E. A. 1989. Comments on genomic genera in the Triticeae (Poaceae). American Journal of Botany 76: 796–805.

Kellogg, E. A., and R. Appels. 1995. Intraspecific and interspecific variation in 5S RNA genes are decoupled in diploid wheat relatives. Genetics 140: 325–343.

Kellogg, E. A., R. Appels, and R. J. Mason-Gamer. 1996. When gene trees tell different stories: the diploid genera of Triticeae (Gramineae). Systematic Botany 21: 321–347.

Kent, W. J. 2002. BLAT—the BLAST-like alignment tool. Genome Research 12: 656–664.

Lanave, C., G. Preparata, C. Sacone, and G. Serio. 1984. A new method for calculating evolutionary substitution rates. Journal of Molecular Evolution 20: 86–93.

Li, L.-F., B. Liu, K. M. Olsen, and J. F. Wendel. 2015. A re-evaluation of the homoploid hybrid origin of *Aegilops tauschii*, the donor of the wheat D-subgenome. New Phytologist 208: 4–8.

Li, W., and A. Godzik. 2006. Cd-hit: a fast program for clustering and comparing large sets of protein or nucleotide sequences. Bioinformatics 22: 1658–1659.

Ling, H. Q., S. C. Zhao, D. C. Liu, J. Y. Wang, H. Sun, C. Zhang, H. J. Fan, et al. 2013. Draft genome of the wheat A-genome progenitor *Triticum urartu*. Nature 496: 87–90.

LöVe, Á. 1984. Conspectus of the Triticeae. Feddes Repertorium 95: 425–521.

Marcussen, T., S. R. Sandve, L. Heier, M. Spannagl, M. Pfeifer, K. S. Jakobsen, B. B. H. Wulff, et al. 2014. Ancient hybridizations among the ancestral genomes of bread wheat. Science 345: 1250092.

Martis, M. M., R. Zhou, G. Haseneyer, T. Schmutzer, J. VráNa, M. KubaláKová, S. KöNig, et al. 2013. Reticulate evolution of the rye genome The Plant Cell 25: 3685–3698.

Mascher, M., H. Gundlach, A. Himmelbach, S. Beier, S. O. Twardziok, T. Wicker, V. Radchuk, et al. 2017. A chromosome conformation capture ordered sequence of the barley genome. Nature 544: 427–433.

Mason-Gamer, R. J. 2001. Origin of North American species of *Elymus* (Poaceae: Triticeae) allotetraploids based on granule-bound starch synthase gene sequences. Systematic Botany 26: 757–768.

Mason-Gamer, R. J. 2005. The b-amylase genes of grasses and a phylogenetic analysis of the Triticeae (Poaceae). American Journal of Botany 92: 1045–1058.

Mason-Gamer, R. J. 2013. Phylogeny of a genomically-diverse group of *Elymus* (Poaceae) allopolyploids reveals multiple levels of reticulation. PLoS ONE 8: e78449.

Mason-Gamer, R. J., and E. A. Kellogg. 1996a. Testing for phylogenetic conflict among molecular data sets in the tribe Triticeae (Gramineae). Systematic Biology 45: 524–545.

Mason-Gamer, R. J., and E. A. Kellogg. 1996b. Chloroplast DNA analysis of the monogenomic Triticeae: phylogenetic implications and genome-specific markers. *In* P. P. Jauhar [ed.], Methods of Genome Analysis in Plants, 301–325. CRC Press, Boca Raton.

Mason-Gamer, R. J., C. F. Weil, and E. A. Kellogg. 1998. Granule-bound starch synthase: structure, function, and phylogenetic utility. Molecular Biology and Evolution 15: 1658– 1673.

Mason-Gamer, R. J., N. L. Orme, and C. M. Anderson. 2002. Phylogenetic analysis of North American *Elymus* and the monogenomic Triticeae (Poaceae) using three chloroplast DNA data sets. Genome 45: 991–1002.

Matsumoto, T., T. Tanaka, H. Sakai, N. Amano, H. Kanamori, K. Kurita, A. Kikuta, et al. 2011. Comprehensive sequence analysis of 24,783 barley full-length cDNAs derived from 12 clone libraries. Plant Physiology 156: 20–28.

Naranjo, T. 2019. The effect of chromosome structure upon meiotic homologous and homoeologous recombinations in Triticeae. Agronomy 9: 552.

Olmstead, R. G., and A. M. Bedoya. 2019. Whole genomes: the holy grail. A commentary on: ‘Molecular phylogenomics of the tribe Shoreeae (Dipterocarpaceae) using whole plastid genomes’. Annals of Botany 123: iv–v.

Orton, L. M., P. Barberá, M. P. Nissenbaum, P. M. Peterson, A. Quintanar, R. J. Soreng, and M. R. Duvall. 2021. A 313 plastome phylogenomic analysis of Pooideae: Exploring relationships among the largest subfamily of grasses. Molecular Phylogenetics and Evolution 159: 107110.

Pattengale, N. D., M. Alipour, O. R. P. Bininda-Emonds, B. M. E. Moret, and A. Stamatakis. 2010. How many bootstrap replicates are necessary? Journal of Computational Biology 17: 337–354.

Petersen, G., and O. Seberg. 1997. Phylogenetic analysis of the Triticeae (Poaceae) based on *rpo*A sequence data. Molecular Phylogenetics and Evolution 7: 217–230.

Petersen, G., and O. Seberg. 2002. Molecular evolution and phylogenetic application of *DMC1*. Molecular Phylogenetics and Evolution 22: 43–50.

Petersen, G., O. Seberg, M. Yde, and K. Berthelsen. 2006. Phylogenetic relationships of *Triticum* and *Aegilops* and evidence for the origin of the **A**, **B**, and **D** genomes of common wheat (*Triticum aestivum*). Molecular Phylogenetics and Evolution 39: 70–82.

Revell, L. J. 2012. phytools: an R package for phylogenetic comparative biology (and other things). Methods in Ecology and Evolution 3: 217–223.

Saski, C., S.-B. Lee, S. Fjellheim, C. Guda, R. K. Jansen, H. Luo, J. Tomkins, et al. 2007. Complete chloroplast genome sequences of *Hordeum vulgare*, *Sorghum bicolor*, and *Agrostis stolonifera*, and comparative analyses with other grass genomes. Theoretical and Applied Genetics 115: 571–590.

Seberg, O., and G. Petersen. 1998. A critical review of concepts and methods used in classical genome analysis. The Botanical Review 64: 372–417.

Seberg, O., and S. Frederiksen. 2001. A phylogenetic analysis of the monogenomic Triticeae (Poaceae) based on morphology. Botanical Journal of the Linnean Society 136: 75–97.

Seberg, O., and G. Petersen. 2007. Phylogeny of Triticeae (Poaceae) based on three organelle genes, two single-copy nuclear genes, and morphology. Aliso 23: 362–371.

Seberg, O., S. Frederiksen, C. Baden, and I. Linde-Laursen. 1991. *Peridictyon*, a new genus from the Balkan peninsula, and its relationship with *Festucopsis* (Poaceae). Willdenowia 21: 87–104.

Seo, T.-K., H. Kishino, and J. L. Thorne. 2008. Incorporating gene-specific variation when inferring and evaluating optimal evolutionary tree topologies from multilocus sequence data. Proceedings of the National Academy of Sciences of the United States of America 102: 4436–4441.

Soreng, R. J., P. M. Peterson, K. Romaschenko, G. Davidse, F. O. Zuloaga, E. J. Judziewicz, T. S. Filgueiras, et al. 2015. A worldwide phylogenetic classification of the Poaceae (Gramineae). Journal of Systematics and Evolution 53: 117–137.

Stamatakis, A. 2006. Phylogenetic models of rate heterogeneity: a high performance computing perspective, Parallel and Distributed Processing Symposium, 20th International. Ieee, Rhodes Island, Greece.

Stamatakis, A. 2014. RAxML Version 8: a tool for phylogenetic analysis and post-analysis of large phylogenies. Bioinformatics 30: 1312–1313.

Sun, G., and B. Salomon. 2009. Molecular evolution and origin of tetraploid *Elymus* species. Breeding Science 59: 487–491.

Sun, G., and X. Zhang. 2011. Origin of the **H** genome in **StH**-genomic *Elymus* species based on the single-copy nuclear gene *DMC1*. Genome 54: 655–662.

Swearington, J., and C. Bargeron. 2016. Invasive Plant Atlas of the United States. Grasses and Grasslike Plants Website https://www.invasiveplantatlas.org/grass.cfm [accessed May 10 2020].

Tanaka, S., K. Yoshida, K. Sato, and S. Takumi. 2020. Diploid genome differentiation conferred by RNA sequencing-based survey of genome-wide polymorphisms throughout homoeologous loci in *Triticum* and *Aegilops*. BMC Genomics 21: 246.

Vaidya, G., D. J. Lohman, and R. Meier. 2011. SequenceMatrix: concatenation software for the fast assembly of multi-gene datasets with character set and codon information. Cladistics 27: 171–180.

Vogel, K. P., and K. B. Jensen. 2001. Adaptation of perennial Triticeae to the eastern Central Great Plains. Journal of Range Management 54: 674–679.

Wang, R. R.-C., and B.-R. Lu. 2014. Biosystematics and evolutionary relationships of perennial Triticeae species revealed by genomic analysis. Journal of Systematics and Evolution 52: 697–705.

Wang, X., X. Yan, Y. Hu, L. Qin, D. Wang, J. Jia, and Y. Jiao. 2022. A recent burst of gene duplications in Triticeae. Plant Communications 3: 100268.

Weitemier, K., S. C. K. Straub, R. C. Cronn, M. Fishbein, R. Schmickl, A. Mcdonnell, and A. Liston. 2014. Hyb-Seq: combining target enrichment and genome skimming for plant phylogenomics. Applications in Plant Sciences 2: 1400042.

West, J. G., C. L. Mcintyre, and R. Appels. 1988. Evolution and systematic relationships in the Triticeae (Poaceae). Plant Systematics and Evolution 1960: 1–28.

Yang, Z. 1994. Maximum likelihood phylogenetic estimation from DNA sequences with variable rates over sites: approximate methods. Journal of Molecular Evolution 39: 306–314.

Zhang, C., M. Rabiee, E. Sayyari, and S. Mirarab. 2018. ASTRAL-III: polynomial time species tree reconstruction from partially-resolved gene trees. BMC Bioinformatics 19: 153.

Zhang, L., X. Zhu, Y. Zhao, J. Guo, T. Zhang, W. Huang, J. Huang, et al. 2022. Phylotranscriptomics resolves the phylogeny of Pooideae and uncovers factors for their adaptive evolution. Molecular Biology and Evolution 39: msac026.

