## Supplement S2-Capture Success for "The phylogeny of Triticeae Dumort. (Poaceae): resolution and phylogenetic conflict based on a genome-wide selection of nuclear loci"

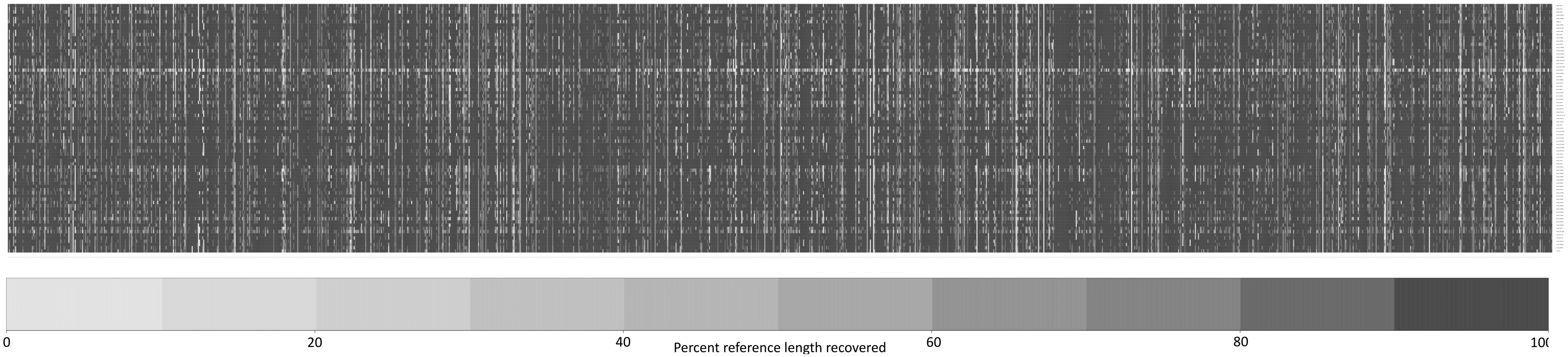


**Appendix S2.** Graphical representation of percent reference coverage for the 1478 *Hordeum* reference sequences (loci) that were used for read-mapping in HybPiper. Each column represents a locus. As explained in the text, 1046 of these were used in subsequent phylogenetic analyses. Each row is a sample, listed from top to bottom in the order shown in Appendix S1. The 21st sample with noticeably poor coverage is an outgroup, *Avena fatua*, which was not included in the phylogenetic analyses.
