## Supplement S4-Chromosome Diagram for "The phylogeny of Triticeae Dumort. (Poaceae): resolution and phylogenetic conflict based on a genome-wide selection of nuclear loci"

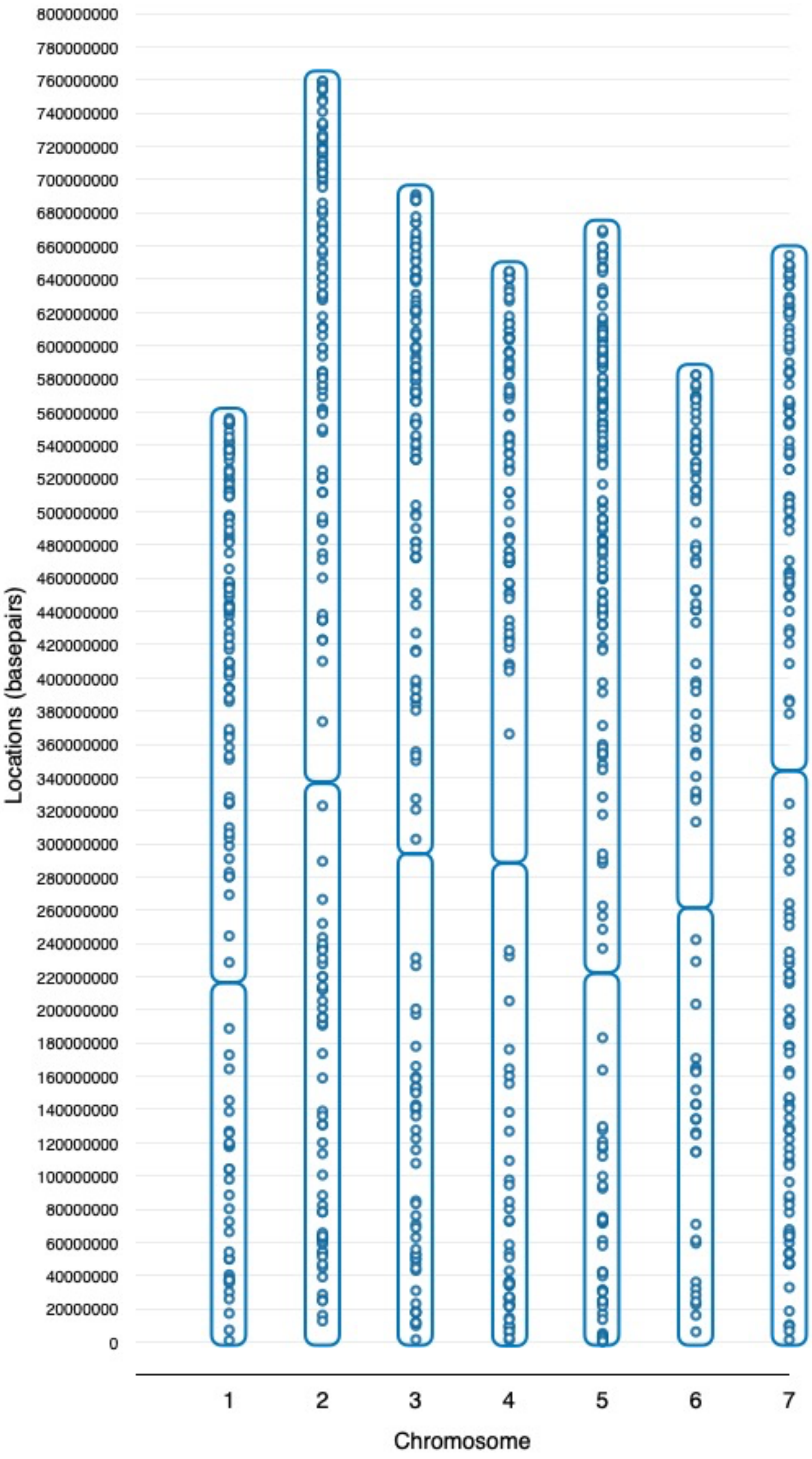


**Appendix S4.** From Appendix S3, approximate positions within the barley genome of the 895 loci used in the concatenated chromosome and chromosome-arm (Appendix S7) analyses, determined using BARLEYMAP (Cantalapiedra et. al. 2015). Rounded rectangles represent chromosome arms; centromere location information is from Mascher et al. (2015).
