## Supplement S5-ASTRAL Tree for "The phylogeny of Triticeae Dumort. (Poaceae): resolution and phylogenetic conflict based on a genome-wide selection of nuclear loci"

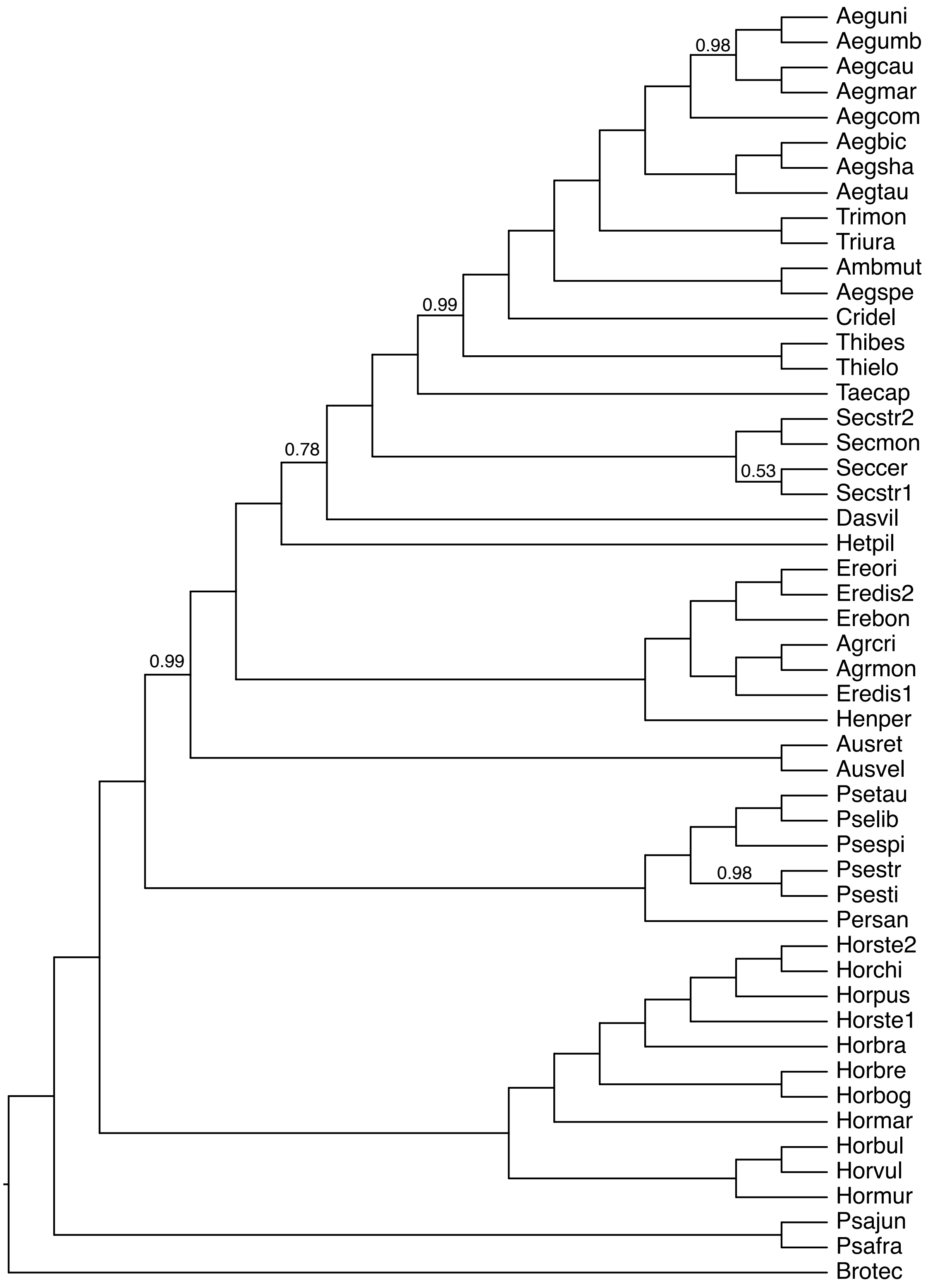


**Appendix S5**. ASTRAL III species tree with 1046 ML gene trees with branches with less than 10% bootstrap support collapsed. Multiple individuals within species were assigned to single terminals, except three that appeared polyphyletic on the ASTRAL III lineage tree of all individuals (not shown): *Eremopyrum distans* (Eredis), *Hordeum stenostachys* (Horste), and *Secale strictum* (Secstr). Posterior probability values less than 1.0 are shown.
