## Supplement S6-Tanglegram for "The phylogeny of Triticeae Dumort. (Poaceae): resolution and phylogenetic conflict based on a genome-wide selection of nuclear loci"

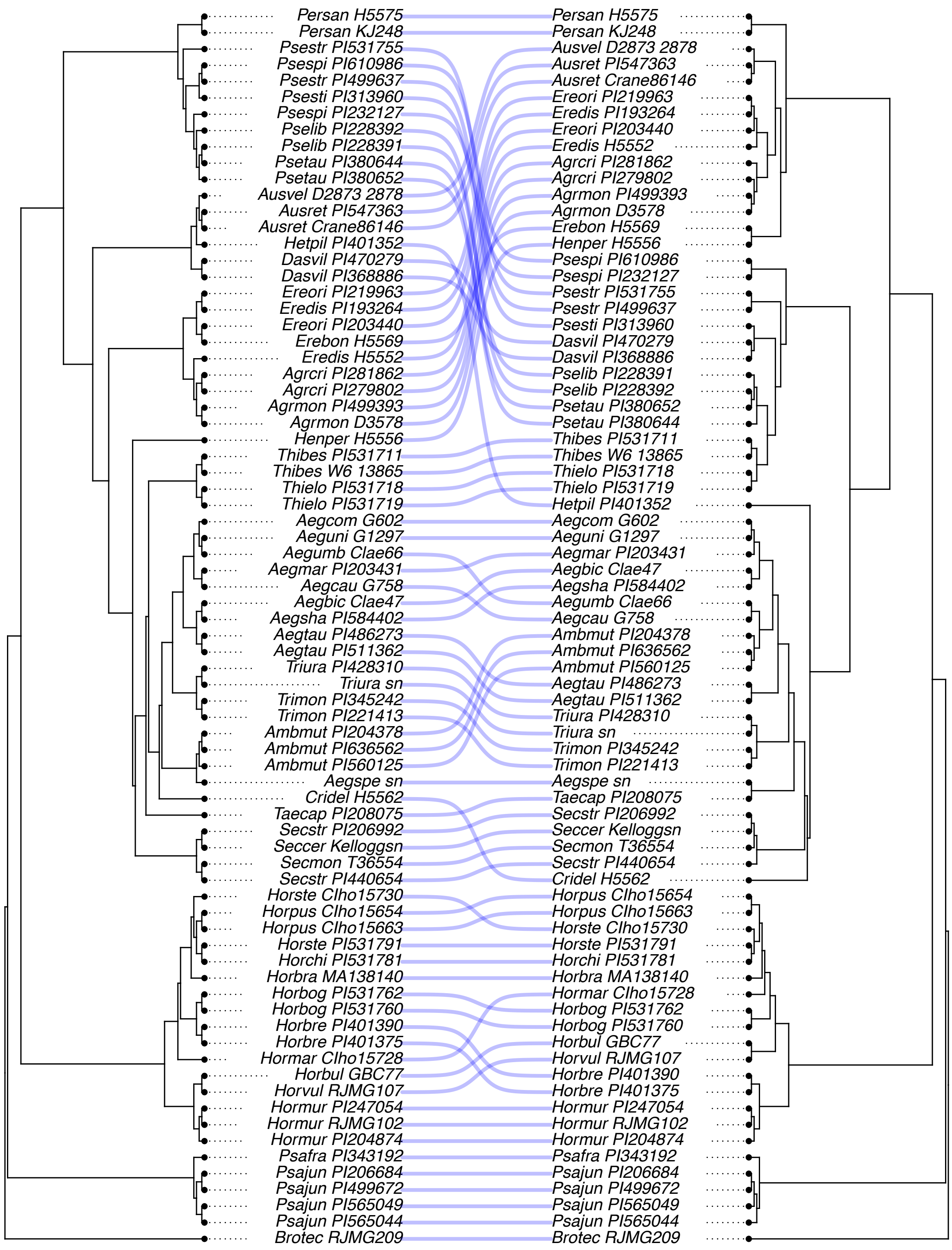


**Appendix S6.** Tanglegram comparison of the cpDNA tree (right) and the concatenated 1046-locus nuclear tree (left) generated using the cophylo function in the R package phytools; some of the differences in topology are discussed in the text.
