## Supplement S7-Chromosome Trees for "The phylogeny of Triticeae Dumort. (Poaceae): resolution and phylogenetic conflict based on a genome-wide selection of nuclear loci"

a) Chromosome 1

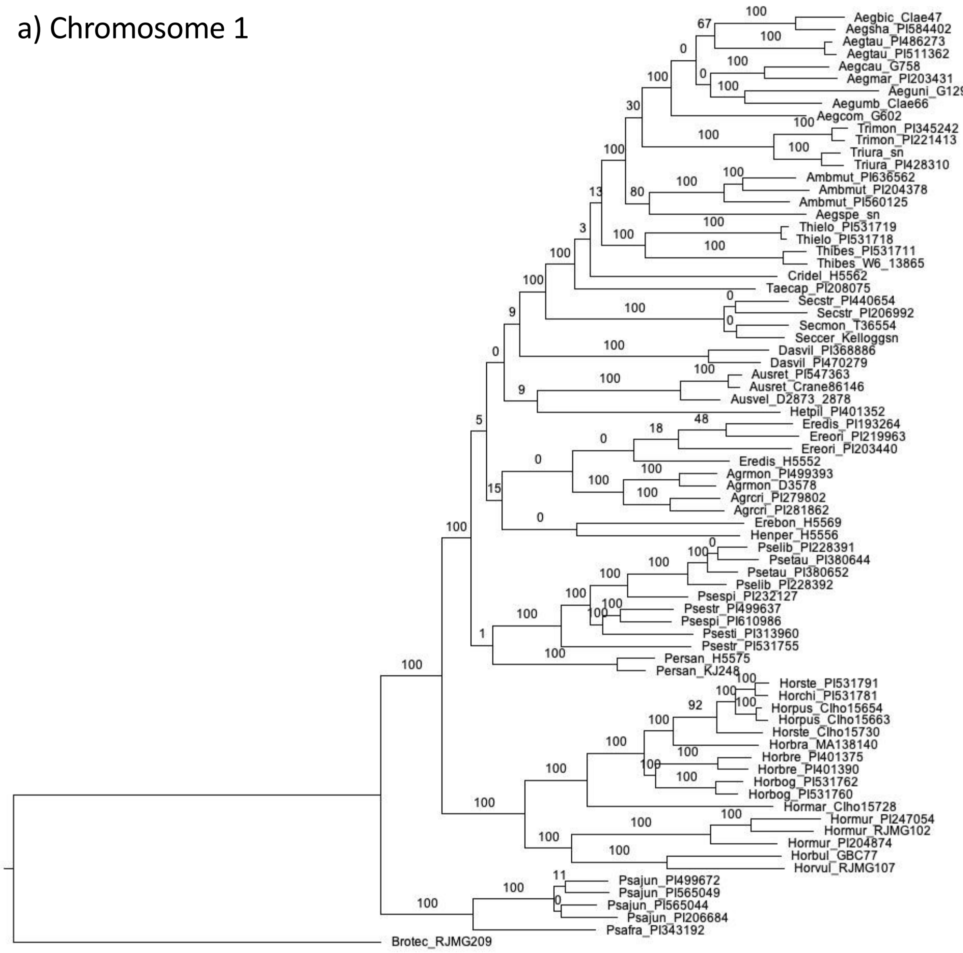


b) Chromosome 1-long

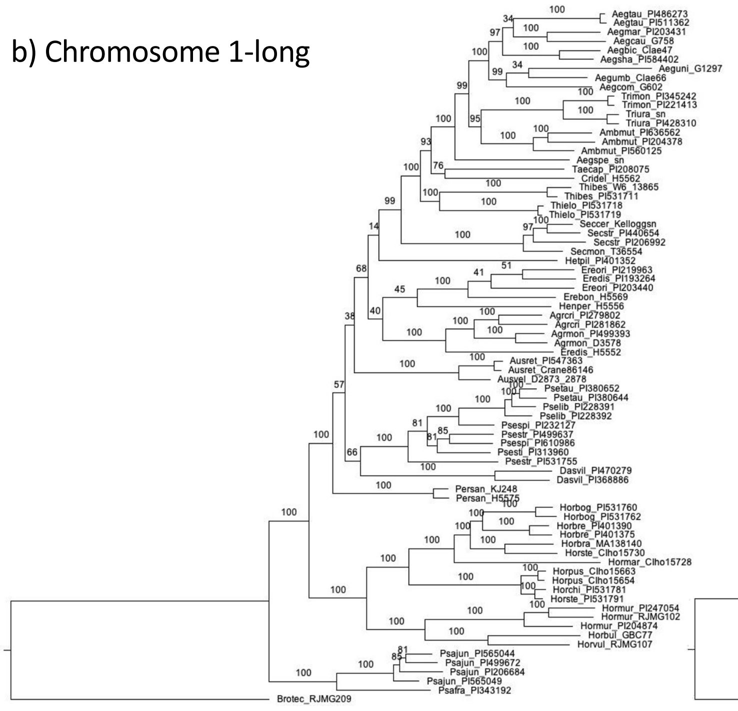

c) Chromosome 1-short

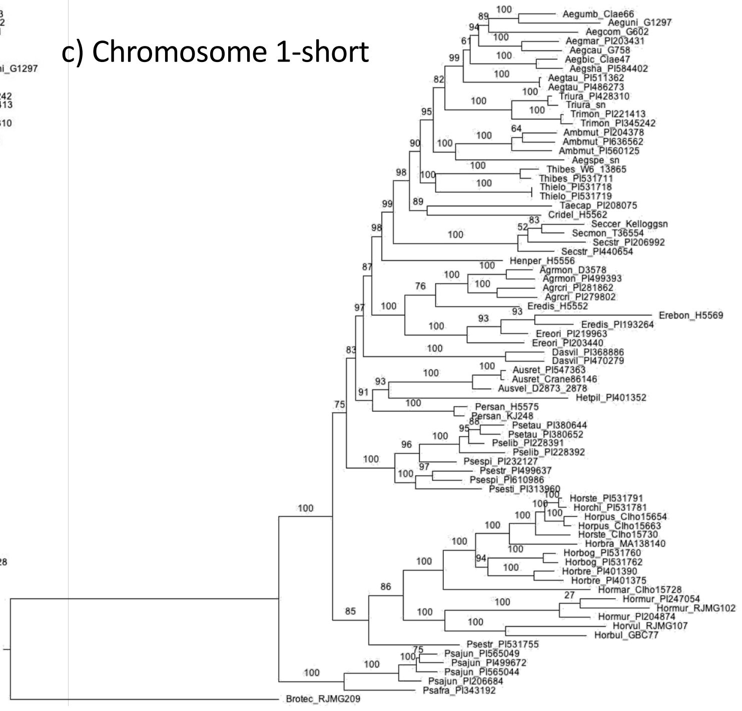

Mason-Gamer and White—Appendix S7

d) Chromosome 2

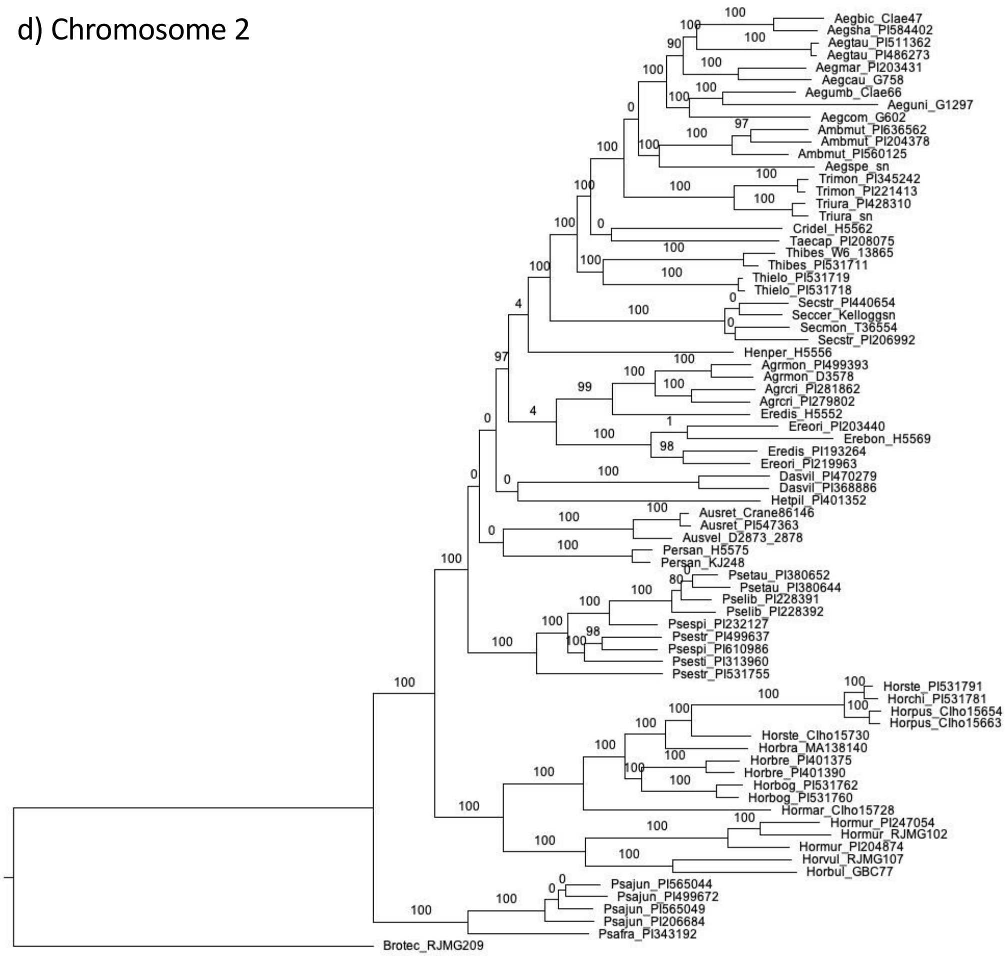


e) Chromosome 2-long

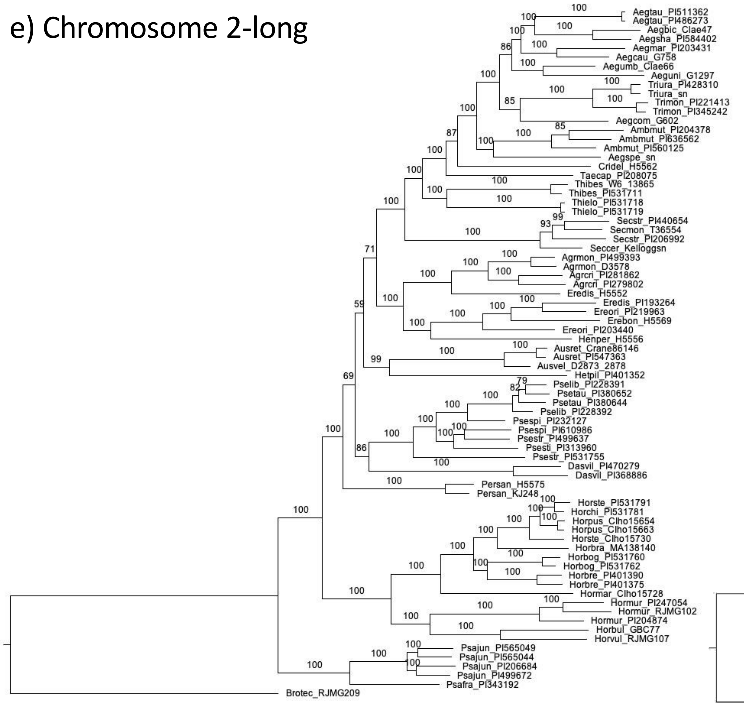


f) Chromosome 2-short

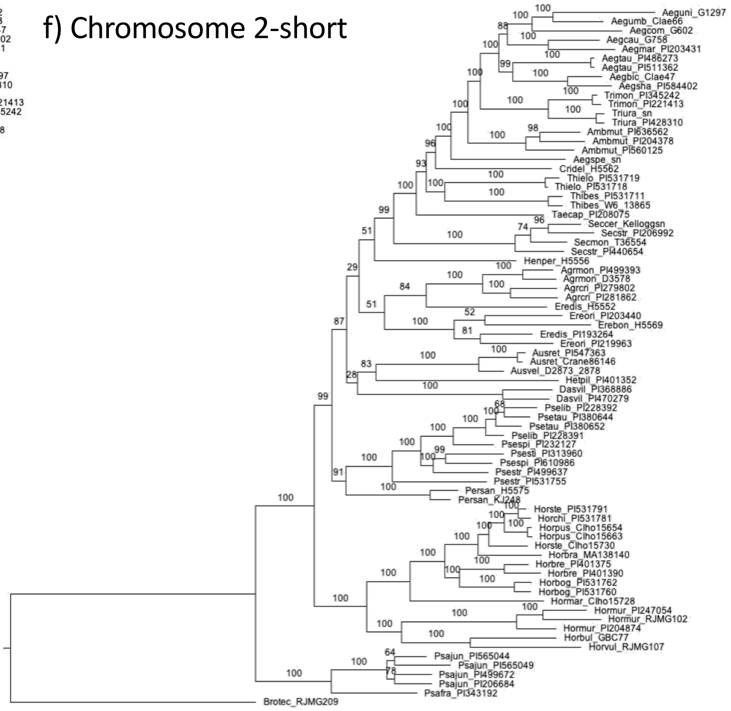

g) Chromosome 3

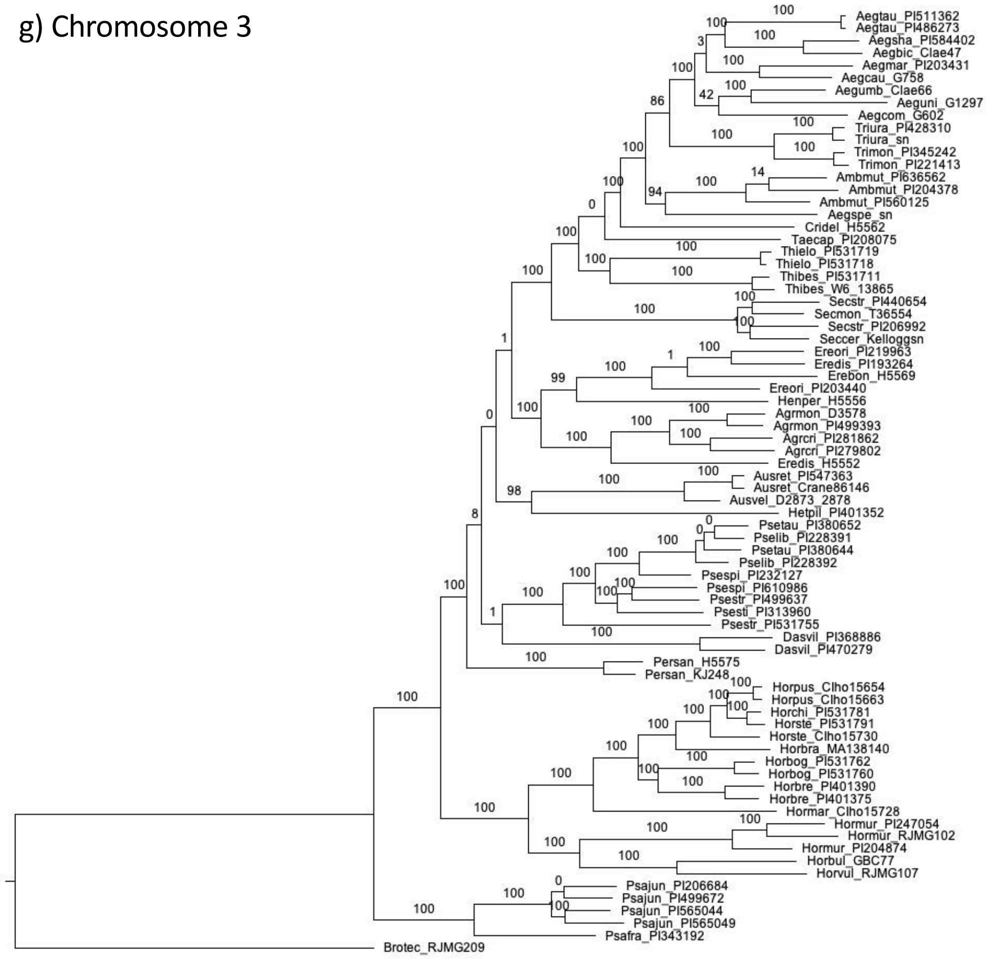


h) Chromosome 3-long

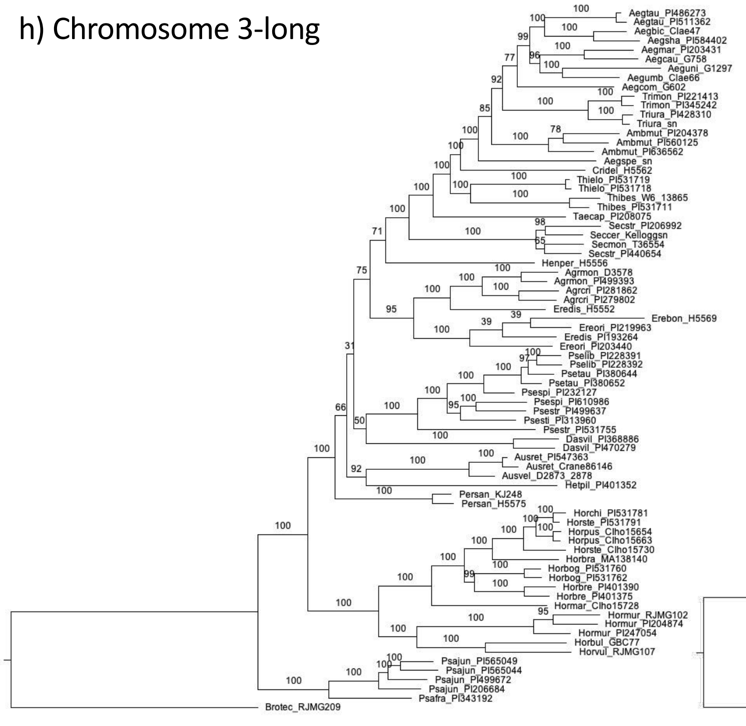

i) Chromosome 3-short

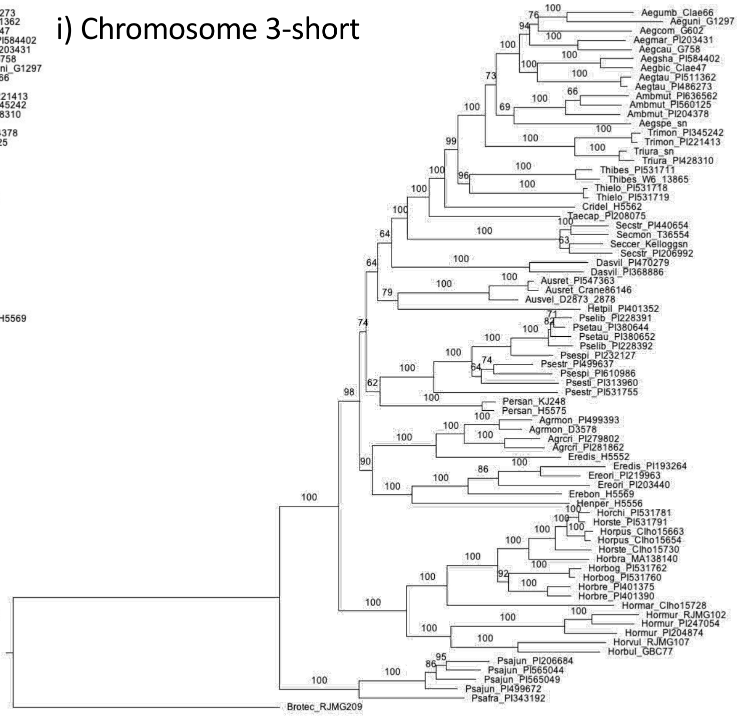

Mason-Gamer and White—Appendix S7

j) Chromosome 4

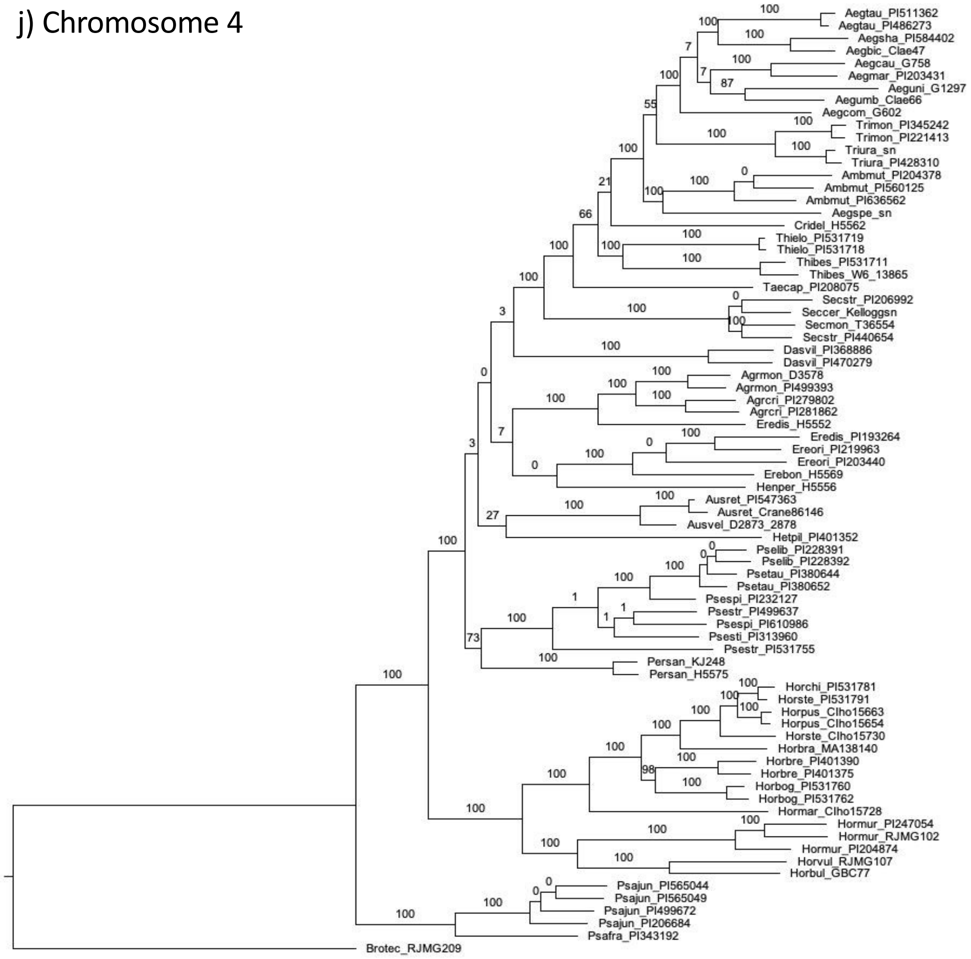


k) Chromosome 4-long

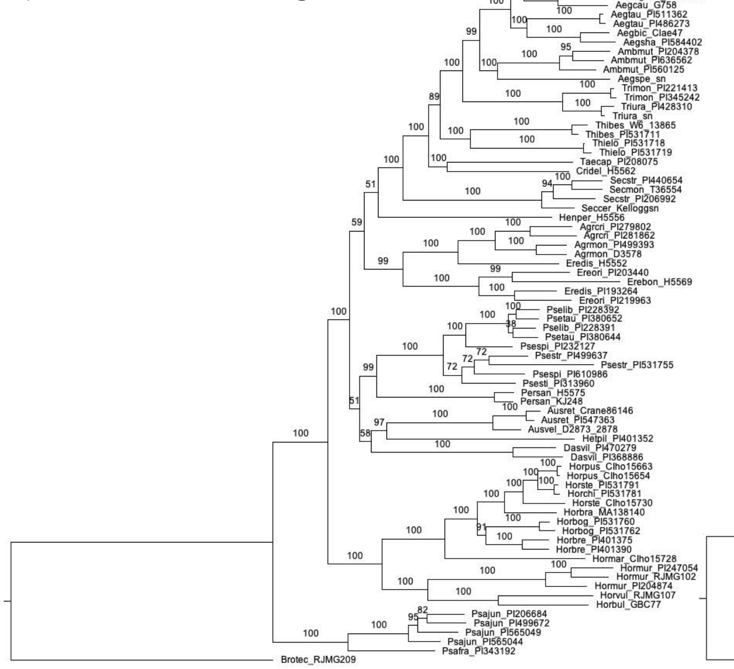


l) Chromosome 4-short

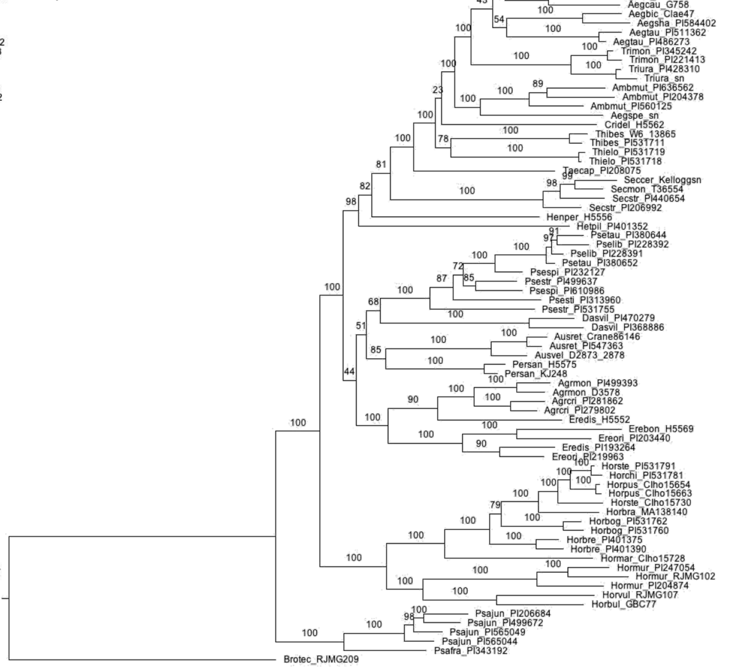

Mason-Gamer and White—Appendix S7

m) Chromosome 5

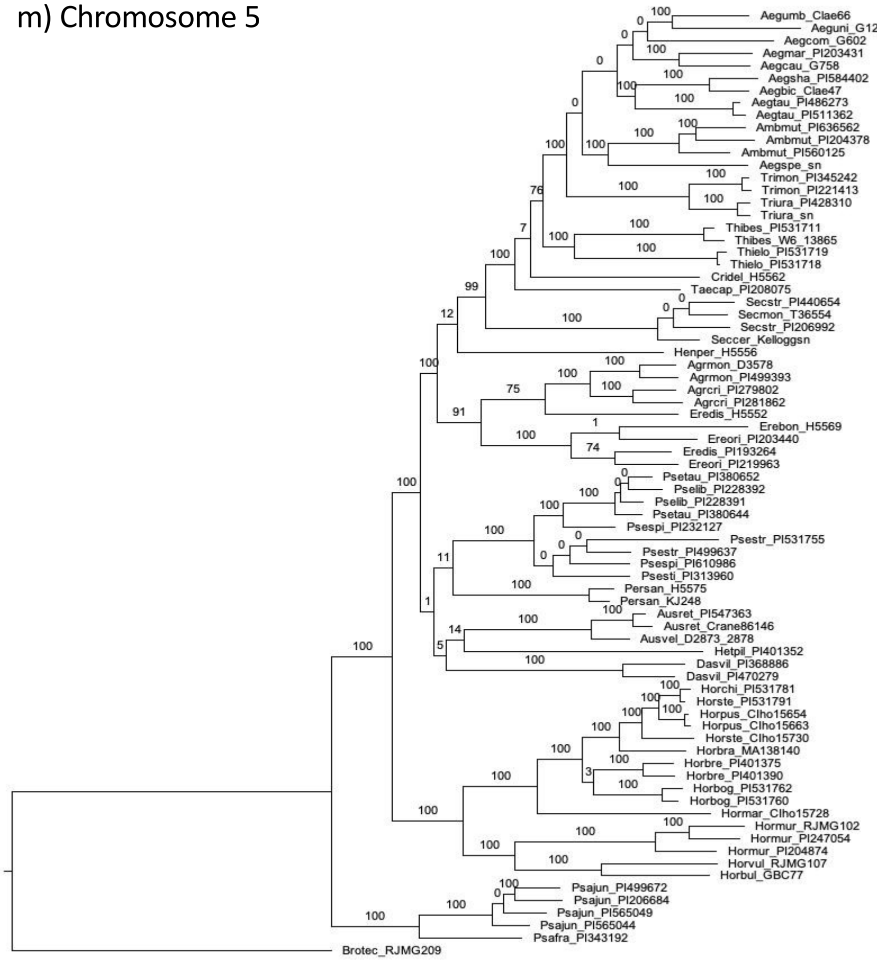


n) Chromosome 5-long

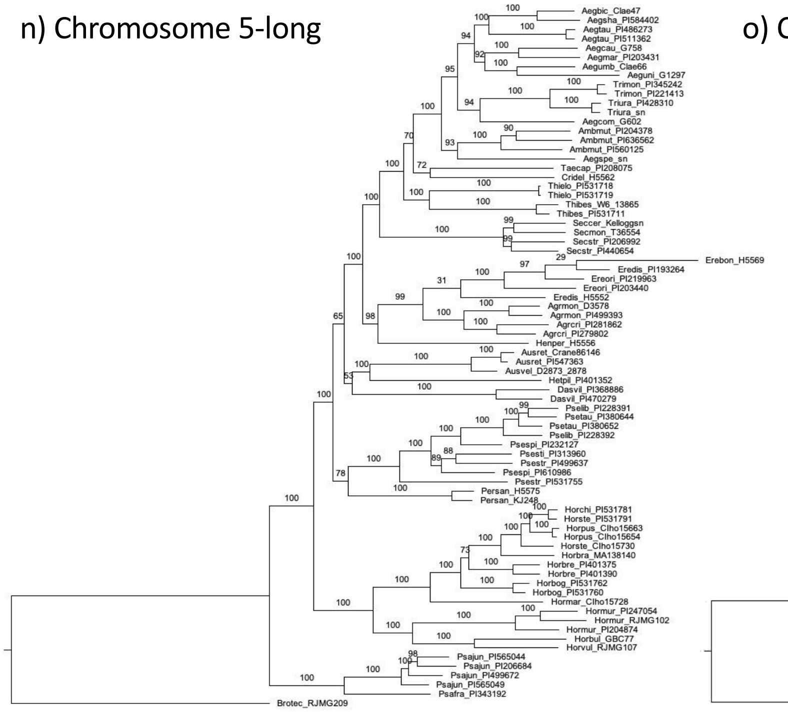

o) Chromosome 5-short

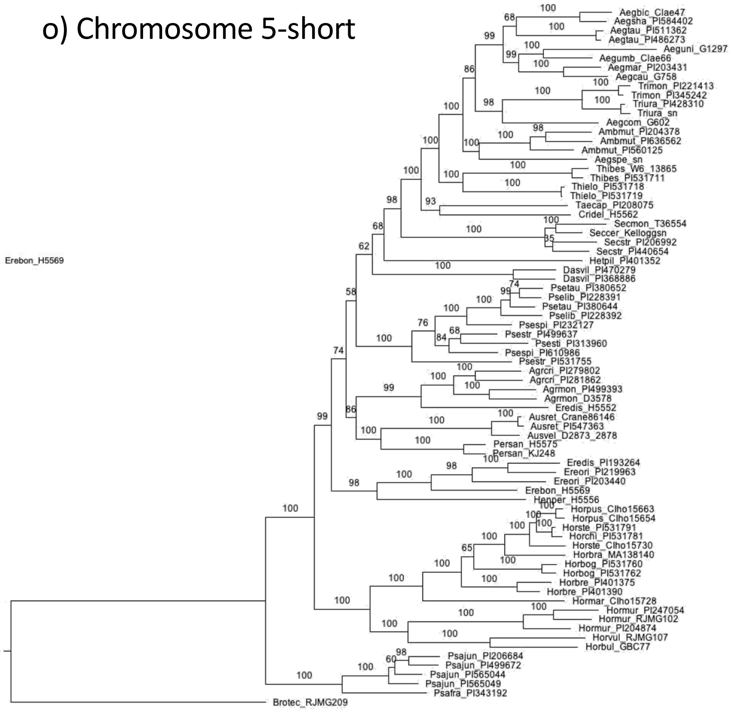

Mason-Gamer and White—Appendix S7

p) Chromosome 6

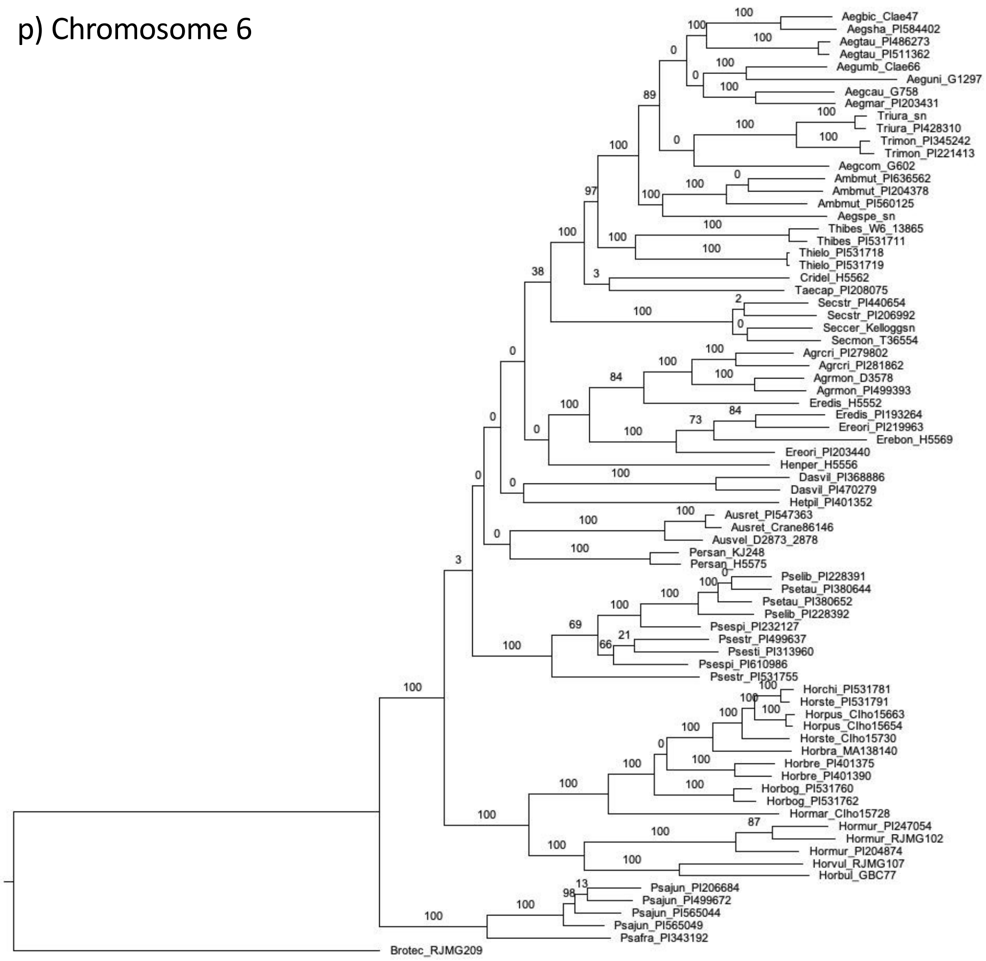


q) Chromosome 6-long

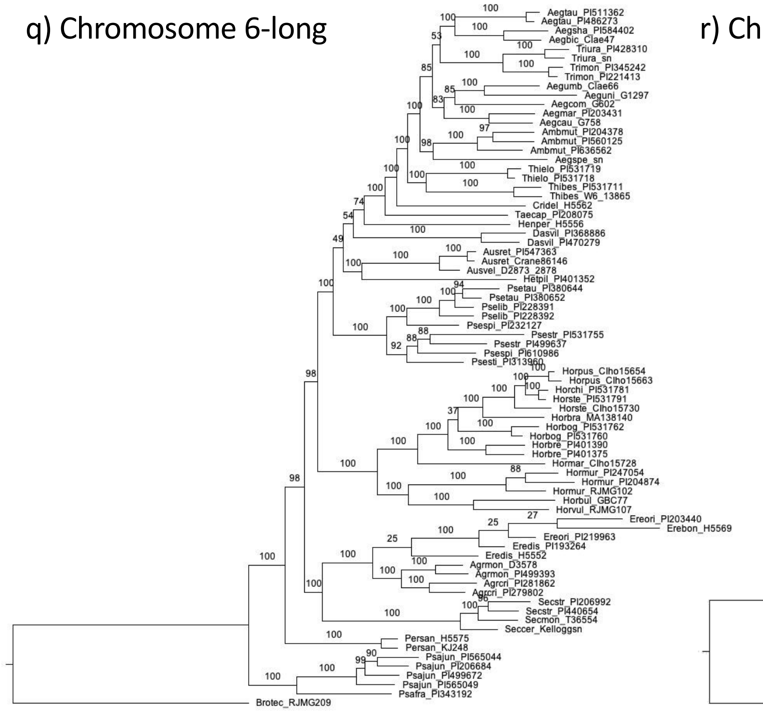


r) Chromosome 6-short

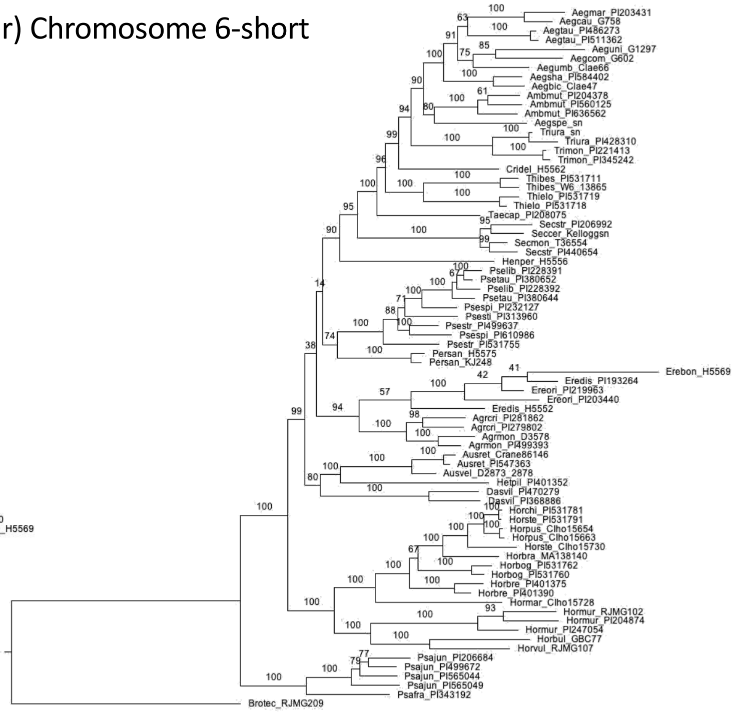

s) Chromosome 7

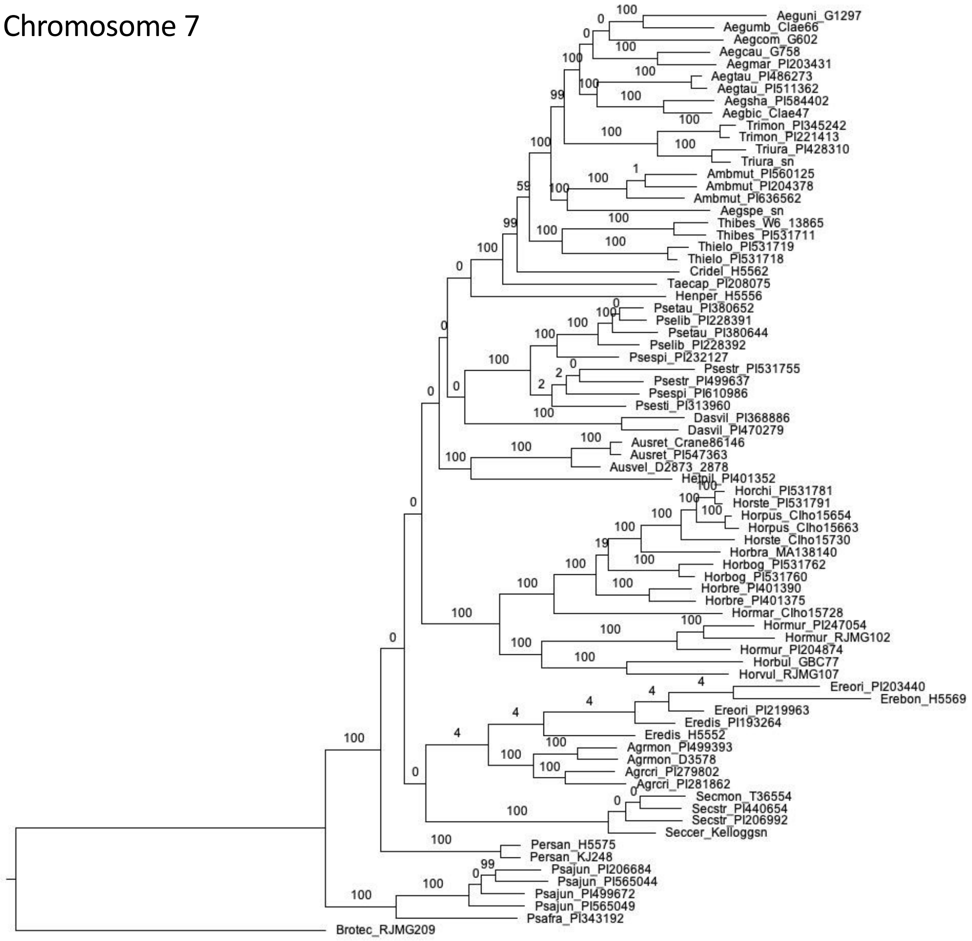


t) Chromosome 7-long

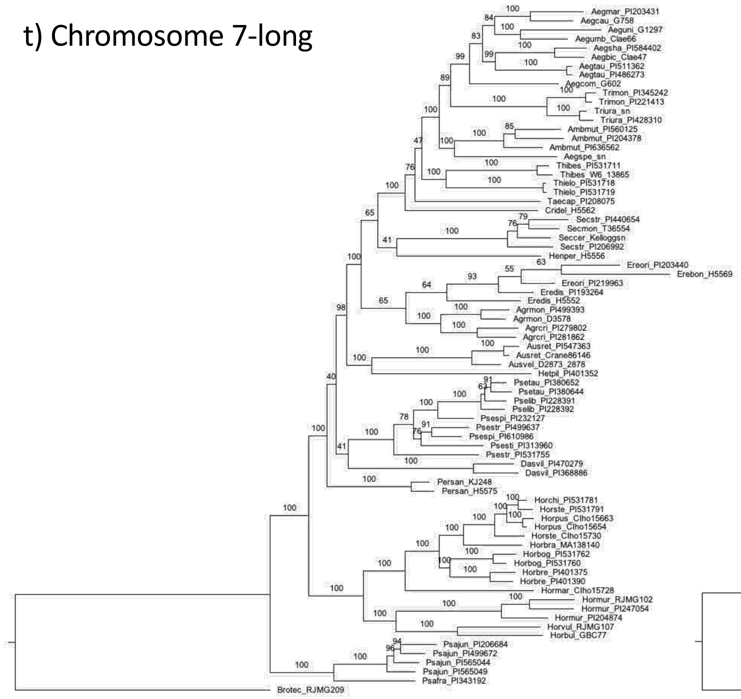

u) Chromosome 7-short

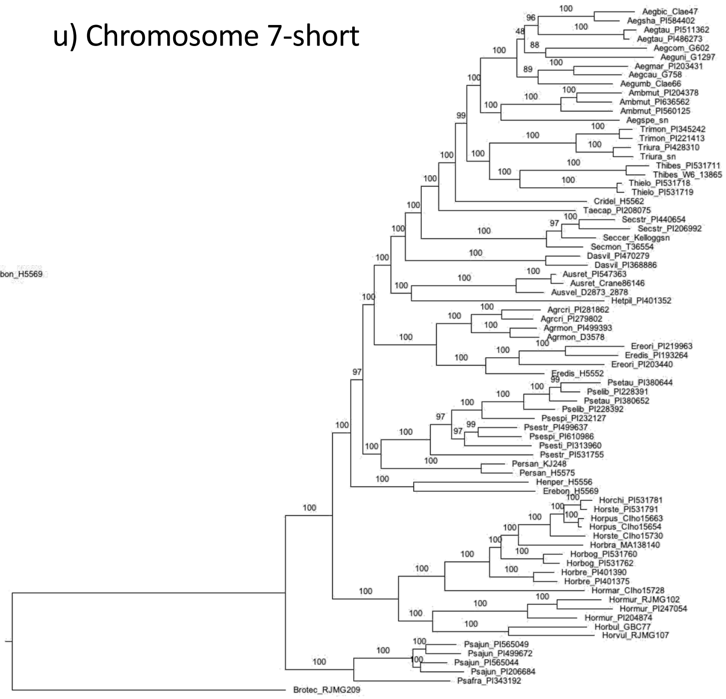

**Appendix S7**, a–u. Individual trees from 21 subsets of loci corresponding to the 7 *Hordeum* chromosomes and 14 chromosome arms (Appendix S3, S4). Each tree represents the best ML tree out of 50 runs under a GTR+gamma model and partitioned by locus, with support from 500 bootstrap replicates under the GTRCAT model. The distinctive tree from the long arm of chromosome 6 (q) is discussed in more detail in the text.
