## Supplement S9-Chromosome 6 Trees for "The phylogeny of Triticeae Dumort. (Poaceae): resolution and phylogenetic conflict based on a genome-wide selection of nuclear loci"

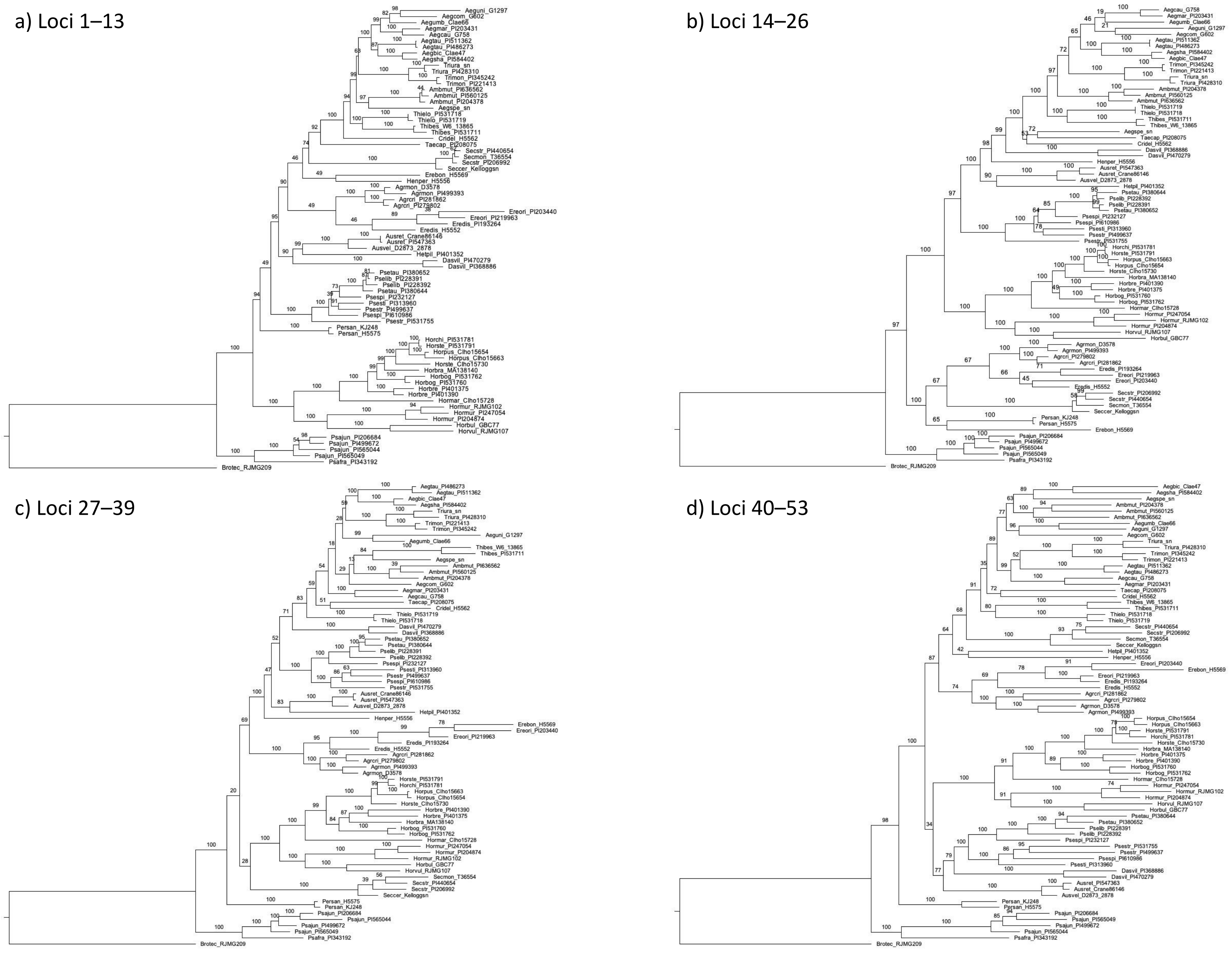


**Appendix S9**, a–d. Maximum-likelihood analyses of four subsets of loci mapped to the long arm of *Hordeum* chromosome 6 (Appendix S8).
